# The Dual Molecular Identity of Vestibular Kinocilia: Bridging Structural and Functional Traits of Primary and Motile Cilia

**DOI:** 10.1101/2025.04.16.647417

**Authors:** Zhenhang Xu, Amirrasoul Tavakoli, Samadhi Kulasooriya, Huizhan Liu, Shu Tu, Celia Bloom, Yi Li, Tirone D. Johnson, Jian Zuo, Litao Tao, Bechara Kachar, David Z. He

## Abstract

Vestibular hair cells (HCs) convert gravitational and head motion cues into neural signals through mechanotransduction, mediated by the hair bundle—a mechanically integrated organelle composed of stereocilia and a kinocilium. The kinocilium, a specialized form of primary cilium, remains incompletely defined in structure, molecular composition, and function. To elucidate its characteristics, we conducted single-cell RNA sequencing of adult vestibular and cochlear HCs, uncovering a selective enrichment of primary and motile cilia–associated genes in vestibular HCs, particularly those related to the axonemal repeat complex. This enrichment of orthologous axonemal-related genes was conserved in zebrafish and human vestibular HCs, indicating a shared molecular architecture. Immunostaining validated the expression of key motile cilia markers in vestibular kinocilia. Moreover, live imaging of bullfrog and mouse HCs from crista ampullaris revealed spontaneous kinociliary motion. Together, these findings define the kinocilium as a unique organelle with molecular features of primary and motile cilia and suggest its previously unknown role as an active, force-generating element within the hair bundle.

## INTRODUCTION

The mammalian inner ear contains auditory and vestibular end organs, which detect sound and motion signals respectively. Hair cells (HCs), the sensory receptors found in both these structures, transduce mechanical stimuli in the form of sound or head movement into electrical signals (Fettiplace, 2017; Gillespie and Müller, 2009). The HCs of the auditory sensory epithelium in the cochlea are categorized as inner and outer HCs (IHCs and OHCs) (Dallos, 1992). The vestibular sensory epithelium lining two otolith organs (utricular and saccular maculae) and three cristae associated with semicircular canals also contains two types of HCs—type I and type II—based on morphology, physiology, and innervation (Eatock et al., 1998; Eatock and Songer, 2011). Although mechanotransduction is a shared feature of all HCs, mammalian cochlear and vestibular HCs differ in morphology and function. One key difference lies in the structure of the hair bundle, which harbors specialized machinery for mechanotransduction. The hair bundle of adult vestibular HCs is composed of actin-rich stereocilia connected via lateral links to a single microtubule-based kinocilium, meanwhile cochlear HCs lose their kinocilia during HC maturation (Leibovici et al., 2005). Kinocilia are highly conserved in non-mammalian vertebrates (Leibovici et al., 2005) and play a pivotal role in establishing hair bundle polarity and mediating Hedgehog and WNT signaling during HC development (Moon et al., 2020; Shi et al., 2022). Despite possessing features of motile cilia such as the canonical “9+2” microtubule arrangement, the kinocilium has long been regarded as a specialized primary cilium (Kikuchi et al., 1989; Wang and Zhou, 2021). While the kinocilium contributes to the bundle mechanics (Baird, 1994; Kindt et al., 2012; Spoon and Grant, 2011), the molecular basis and function of this unique organelle in adult vestibular HCs remain unknown.

In the current study we utilized single-cell RNA-sequencing (scRNA-seq) to examine the transcriptomes of 1,522 HCs isolated from cochlear and vestibular sensory epithelia of adult CBA/J mice Comparisons of the molecular profiles of the four HC types identified novel marker genes as well as shared and unique genes associated with mechanotransduction, ion channels, and pre- and post-synaptic structures. Notably, our analysis revealed a significant enrichment of genes related to primary and motile cilia in vestibular HCs, particularly those linked to the 96-nm axonemal repeat complex, a hallmark feature of motile cilia. Orthologous axonemal-related genes were also detected in zebrafish HCs and human vestibular HCs. We utilized transmission electron microscopy (TEM) to examine the ultrastructure of kinocilia and immunostaining to detect expression of key motile cilia proteins in vestibular HCs. We also used live imaging to examine kinocilia motion in bullfrog and mouse crista ampullaris. We modeled the atomic architecture of the 96-nm repeat, the core framework of the kinocilium axoneme. Together, these findings establish the kinocilium as a distinct organelle with molecular hallmarks of motile cilia and suggest it functions as an active, force-generating hair bundle component, influencing the mechanosensitivity of the kinocilium-bearing HCs across nonmammalian and mammalian species. In addition, our transcriptomic analysis provides new insight into the molecular mechanisms underlying phenotypical differences among the four different HC types in the adult mouse inner ear.

## RESULTS

Solitary cells were isolated from the whole basilar membrane (together with the organ of Corti, Fig. 1A) and vestibular end organs from 10-week-old CBA/J mice. Cells isolated from the cochlea include IHCs, OHCs, supporting cells (SCs), spiral ganglion neurons (SGNs), and other accessory cells. Some examples of individual IHCs and OHCs are shown in Fig. 1A. Cells isolated from vestibular sensory epithelia include type I HCs, type II HCs, SCs, vestibular neurons and other cell types. Vestibular HCs can be identified based on morphological features: Type I HCs are flask-shaped with a narrow neck while type II HCs are cylindrical and short (Burns and Stone, 2017; Eatock et al., 1998; Lysakowski and Goldberg, 1997; Pujol et al., 2014; Ricci et al., 1997). Type II HCs and SCs also express *Sox2*/SOX2 (Fig. 1B), which is a marker for these cell types in the vestibular sensory epithelium (Jan et al., 2021; Wilkerson et al., 2021). Some representative images of type I and type II HCs isolated from maculae of utricle and saccule as well as from crista ampullaris are presented in Fig. 1C.

**Fig. 1:**
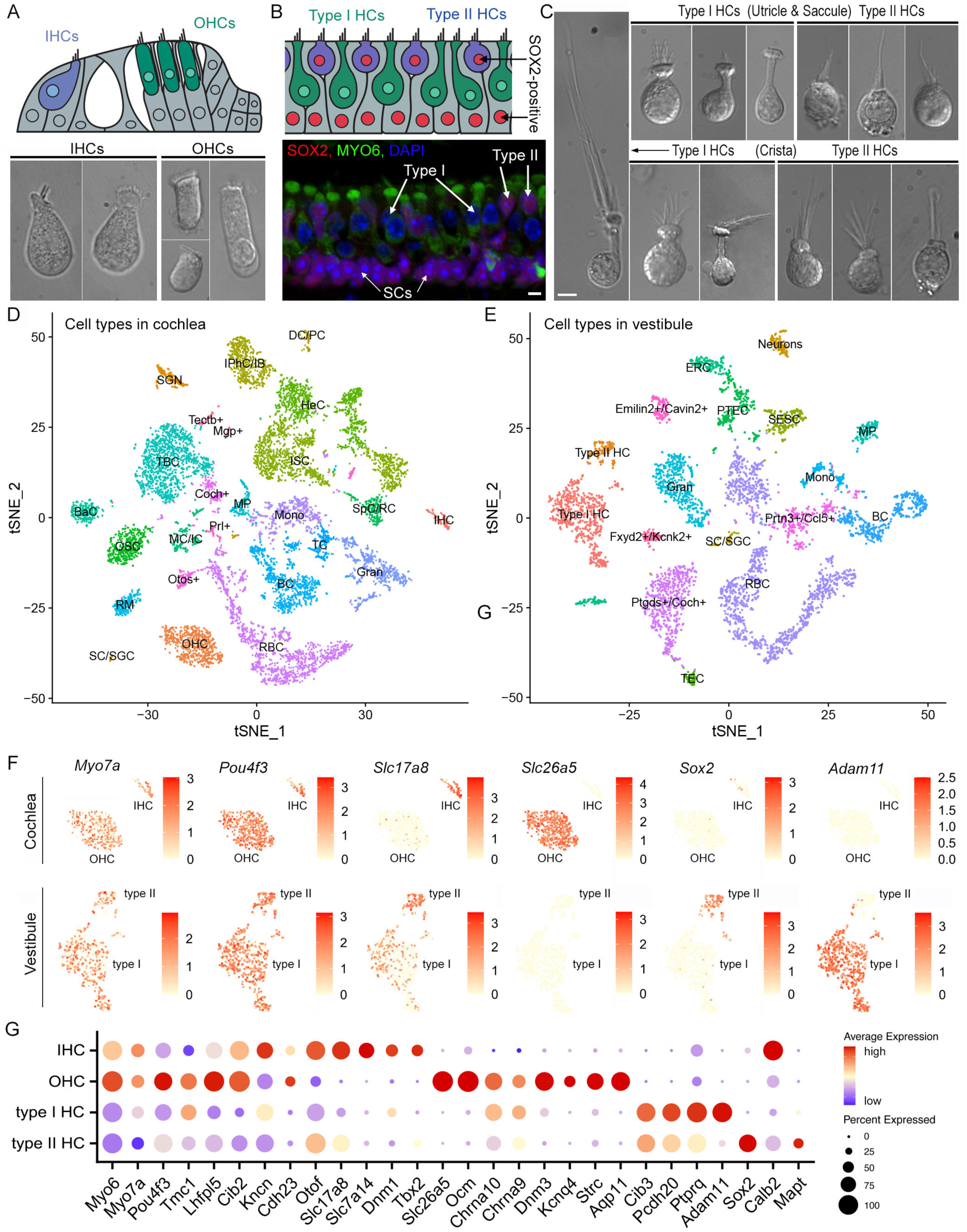
Single-cell transcriptional atlas of cochlear and vestibular cells. **A**: Schematic drawing of the organ of Corti (top panel) and representative images of IHCs and OHCs from adult mouse cochleae. **B**: Schematic drawing of the utricle (top panel) and confocal images of the utricle prepared from an adult mouse. HCs are stained with MYO6, SOX2 and DAPI. SOX2-positive cells include type II HCs and SCs underneath HCs. **C**: Representative images of type I and type II HCs from utricular and saccular maculae as well as crista ampullaris from adult mice. **D**, **E**: tSNE plots of distinct cell types detected in the adult CBA mouse cochlea (D) and utricle, saccule and crista. Different cell types are color-coded. **F**: Feature plots of the expression 6 marker genes in different HC populations. **G**: Dot plot heatmap of average expression and cellular detection rate of 28 representative marker genes in different HC types in cochleae and vestibular end organs. Abbreviations: IHC (inner HC); OHC (outer HC); SGN (spiral ganglion neuron); SC (Schwann cell)/SGC (satellite glial cell); DC (Deiters’ cell)/PC (pillar cell); IPhC (inner phalangeal cell)/IB (inner border cell); ISC (inner sulcus cell); HeC (Hensen’s cell); SpC (spindle cell)/RC (root cell); MC (marginal cell)/IC (intermediate cell); BaC (basal cell); MP (macrophage); RM (Reissner’s membrane); BC (B cell); TC (T cell); Gran (granulocyte); Mono (monocyte); RBC (red blood cell); OSC (outer sulcus cell); TBC (tympanic border cell). Type I HC (type I HC); Type II HC (type II HC); SESC (sensory epithelial SC); TEC (transitional epithelial cell); PTEC (peripheral transitional epithelial cell); ERC (epithelial roof cell). Neurons (vestibular neurons). For those cells whose definite identities cannot be annotated, the top expressed genes were used for identification and annotation. These cells include Tectb^+^, Mgp^+^, Coch^+^, Prl^+^, Otos^+^, Emilin2^+^/Cavin2^+^, Fxyd2^+^/Kcnk2^+^, Prtn3^+^/Ccl5^+^, and Ptgds^+^/Coch^+^.

To comprehensively assess the molecular profiles of inner ear HCs, we conducted scRNA-seq using cells isolated from the auditory and vestibular sensory epithelia. The within-tissue cellular diversity and identity in the cochlear and vestibular sensory epithelia were assessed by a t-distributed Stochastic Neighbor Embedding (t-SNE) analysis followed by clustering and annotations of cell types based on the expression of known marker genes (Fig. 1D,E)(Burns et al., 2015; Jan et al., 2021; McInturff et al., 2018; Ranum et al., 2019; Wilkerson et al., 2021). Upon cluster annotation and normalization across biological repeats, HC types were separated for the downstream analysis by their known marker genes shown in feature plots (Fig. 1F) and dot plots (Fig. 1G). A total of 1,522 individual cells were identified as HCs, including 131 IHCs, 668 OHCs, 588 type I HCs, and 135 type II HCs for our downstream analysis. We aggregated the number of reads for each gene across all single cells to generate pseudo-bulk expression profiles for different HC types (Table S1).

### 1. Similarities among HC types

We utilized the pseudo-bulk and single-cell gene expression profiles to compare the four HC types. Since adult IHC and OHC transcriptomes have been compared extensively in the past (Li et al., 2018; Ranum et al., 2019), we focused our analyses on the differences between cochlear and vestibular HCs, as well as between type I and type II HCs. Principal component analysis (PCA) was used to denote variance across the four types of HCs. The first principal component (PC1) shows that the most dramatic differences are tissue-based—between cochlear and vestibular HCs (Fig. 2A). The distribution across PC2 indicates that the similarity between type I and type II HCs is greater than the similarity between IHCs and OHCs, suggesting more homogeneity among vestibular HCs. Gene expression profiles of single HCs were also used to analyze similarities among the four different HC types (Fig. 2B). Despite heterogeneity found among individual HCs of each type, three-dimensional PCA visualization of single cells draws similar conclusions to the pseudo-bulk PCA, suggesting higher transcriptomic similarity between vestibular HCs in contrast to cochlear HCs.

**Figure 2:**
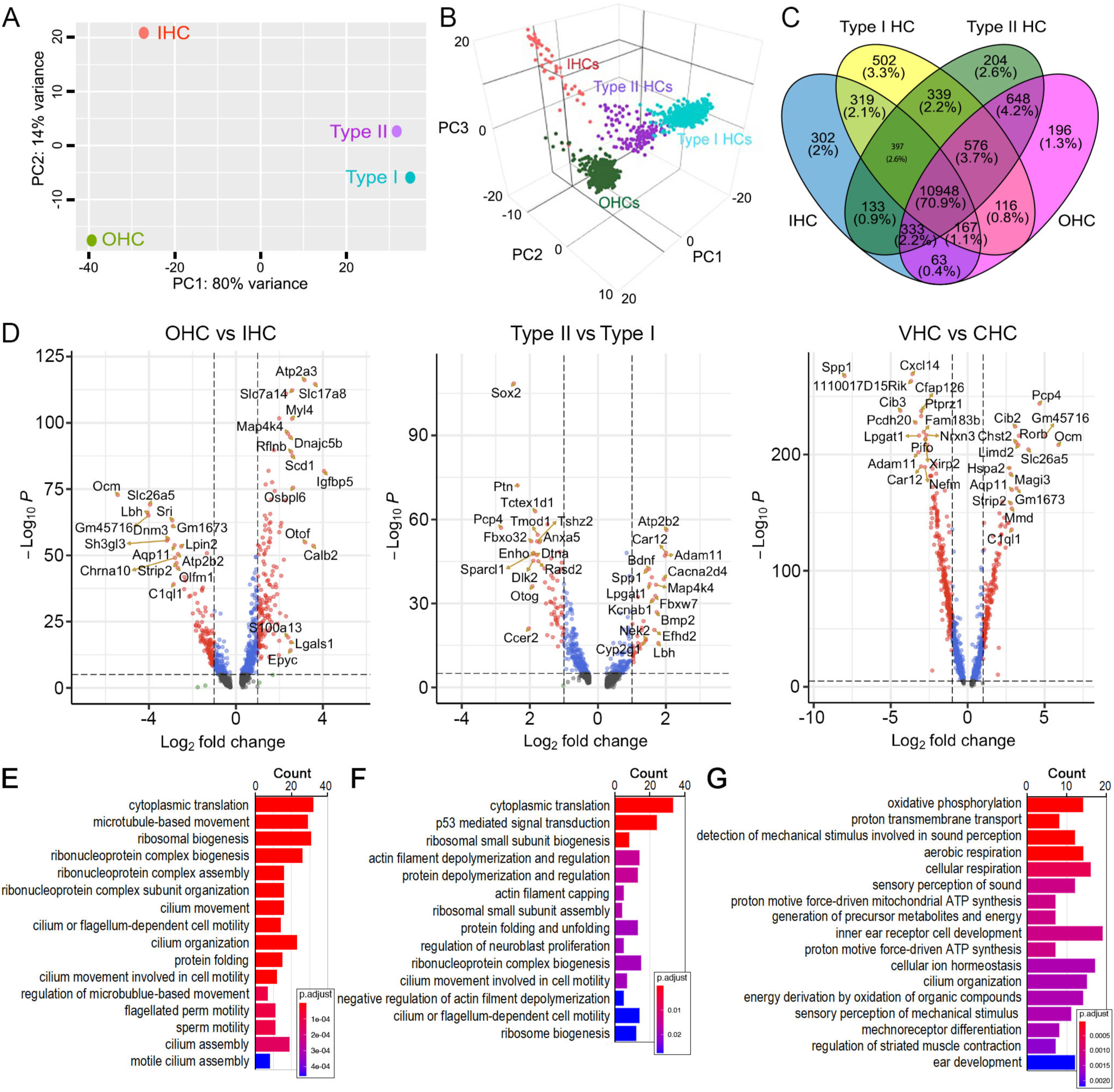
Similarity and difference among different HC types and biological processes enriched in cochlear and vestibular HCs. **A**: PCA plot showing similarity based on pseudo bulk RNA-seq data from four HC types. **B**: PCA plot showing similarity based on individual HC gene expression among the four HC types. **C**: Venn diagram depicting the number of expressed genes (RPKM > 0) in four HC types. **D**: Volcano plot showing differentially expressed genes between different HC types. Red dots indicate differentially expressed genes with P-value < 10^e^-5 and log_2_ fold change > 1. Only the top 20 differentially expressed genes are labeled. **E**: Biological processes enriched in vestibular HCs compared to cochlear HCs. Biological processes related to motile cilia are enriched. **F**: Biological processes enriched in type I HCs compared to type II HCs. **G**: Biological processes enriched in type II HCs compared to type I HCs.

### 2. Differentially expressed genes in HC types

Next, we examined the number of genes that are shared and unique among the four types of HCs based on pseudo-bulk expression profiles (Fig. 2C). Although approximately 71% of the detected genes are shared among all four types of HCs, the smaller proportion of unique genes in each HC population may underlie their specific biological identities. We performed pairwise differentially expressed gene (DEG) analyses between within-tissue HC types, as well as between cochlear and vestibular HCs (Fig. 2D). DEGs were defined as those with an expression level above 0 and a minimum of 2-fold change (log2 ≥ 1) between the two cell populations with statistical significance of *p* ≤ 0.01.

DEGs between adult IHCs and OHCs have been analyzed before using cell type-specific microarray and bulk RNA-seq techniques (Li et al., 2018; Liu et al., 2014). Here, our comparison revealed differential enrichment of 154 and 123 genes in IHCs and OHCs respectively (Fig. 2D). We show expression of previously characterized DEGs in IHCs (e.g., *Otof*, *Slc17a8*) and OHCs (e.g., *Ocm*, *Slc26a5*, *Chrna10*) as well as genes whose functions have not yet been characterized, including *Atp2a3*, *Calb2*, *Dnajc5b*, *Ripor3*, and *Scd1* in IHCs, and *Aqp11*, *Dnm3*, and *Sh3gl3* in OHCs. Our DEG analysis comparing IHCs and OHCs is consistent with the previous studies (Li et al., 2018; Liu et al., 2014).

We next compared DEGs between type I and type II HCs (Fig. 2D). Although some new marker genes of adult type I and type II HCs from mouse utricles have been identified (McInturff et al., 2018), differential gene expression analysis between type I and type II HCs has not been conducted extensively. Our analysis revealed enrichment of 35 DEGs in type I HCs and 59 DEGs in type II HCs. Except for a few genes such as *Otog* and *Bmp2*, the roles of these DEGs in vestibular HCs have not been examined.

Grouping the tissue-specific subtypes together, we calculated DEGs between cochlear and vestibular HCs (Fig. 2D). This comparison identified 674 DEGs enriched in cochlear HCs and 887 DEGs enriched in vestibular HCs. We note that many vestibular DEGs (such as *Adam11*, *Car12*, *Nrxn3*, *Lpgat1*, *Spp1*, *Pcdh20*, *Cfap126*) are related to the secretion of extracellular matrix protein, cell-matrix interactions, cell adhesion, mineralized matrix, calcification, acid-base balance, and cilia. Meanwhile, many of the genes enriched in cochlear HCs (such as *Slc25a5*, *Strip2*, *C1ql1*, *Rorb*, and *Ocm*) are related to the extensively studied unique structure and function of cochlear HCs. It is interesting that *Cib2* is differentially expressed in cochlear HCs while *Cib3* is enriched in vestibular HCs. *Cib2* and *Cib3* act as an auxiliary subunit of the sensory mechanoelectrical transduction (MET) channel in HCs (Giese et al., 2017; Riazuddin et al., 2012; Wang et al., 2023) and mutations of these two genes are associated with deafness and Usher syndrome 1J (Riazuddin et al., 2012).

To examine the functional relevance of the calculated DEGs, we conducted overrepresentation analysis (ORA). Since the molecular properties of vestibular HCs are less known compared to cochlear HCs, we looked more closely at the biological processes enriched in vestibular HCs compared to cochlear HCs, which included gene ontology (GO) terms related to cilium organization and microtubule-based cilia motility (highlighted by red asterisks in Fig. 2E). We also conducted ORA between type I and type II HCs. In type I HCs, enriched processes included those related to cytoplasmic translation, p53-mediated signal transduction, actin filament depolymerization and regulation, and cilium or flagellum-dependent cell motility (Fig. 2F). Terms enriched in type II HCs are related to oxidative phosphorylation, proton transmembrane transport, aerobic and cellular respiration, detection of mechanical stimulus involved in sound perception and mechanoreceptor differentiation (Fig. 2G).

### 3. Marker genes for different HC types and genes related to HC specialization

Previous studies have characterized some marker genes in different HC types. We assessed the expression of previously characterized and newly identified genes related to HC structure and function across all four HC types. Many genes are expressed in all four HC types (Fig. 3A). Among these are well-characterized genes such as *Cib2*, *Otof*, *Kcna10*, *Ptprq*, *Tmc1*, and *Espn*, while other genes such as *Tjap1*, which encodes a tight junction-associated protein, are less known. We noted genes that were enriched in a tissue-specific manner. For example, *Cdh23*, *Rorb* and *Osbp2* are more highly expressed in cochlear HCs than in vestibular HCs, while genes such as *Ldhb*, *Fbxo32*, *and Gsn* are enriched in vestibular HCs but only weakly expressed in cochlear HCs. Several genes expressed in vestibular HCs with no expression in cochlear HCs (such as *Cfap43*, *Cfap126*, *Cib3*, *Cxcl14*, *Pcdh20*, *Pifo*, *Slc9a3r2*, *Tmc2*) could potentially be used as vestibular HC markers (Fig. 3A). Moreover, *Adam11*, *Cacna2d4*, *Car12*, and *Shank2* are found to be only expressed in type I HCs, while *Ccer2*, *Cfap45*, *Dlk2*, and *Rprm* are only expressed in type II HCs. Our analysis also revealed new marker genes for IHCs (*Atp2a3*, *Rims2*, *Ripor3*) and OHCs (*Aqp11*, *Mmd*, *Sh3gl3*)*. Cfap43*, *Cfap44*, *Cfap45*, *Cfap126*, *Kif3*, *Mlf1* and *Pifo*, enriched in vestibular HCs, are all associated with kinocilium structure and function. We utilized single-molecule fluorescent *in situ* hybridization (smFISH) and immunostaining techniques to validate the expression of 12 genes in HCs (Fig. 3B, C). The expression patterns of these genes and proteins were highly consistent with our observations from scRNA-seq analysis. Overall, our results identified novel marker genes and previously unidentified expression patterns among the four HC types.

**Figure 3:**
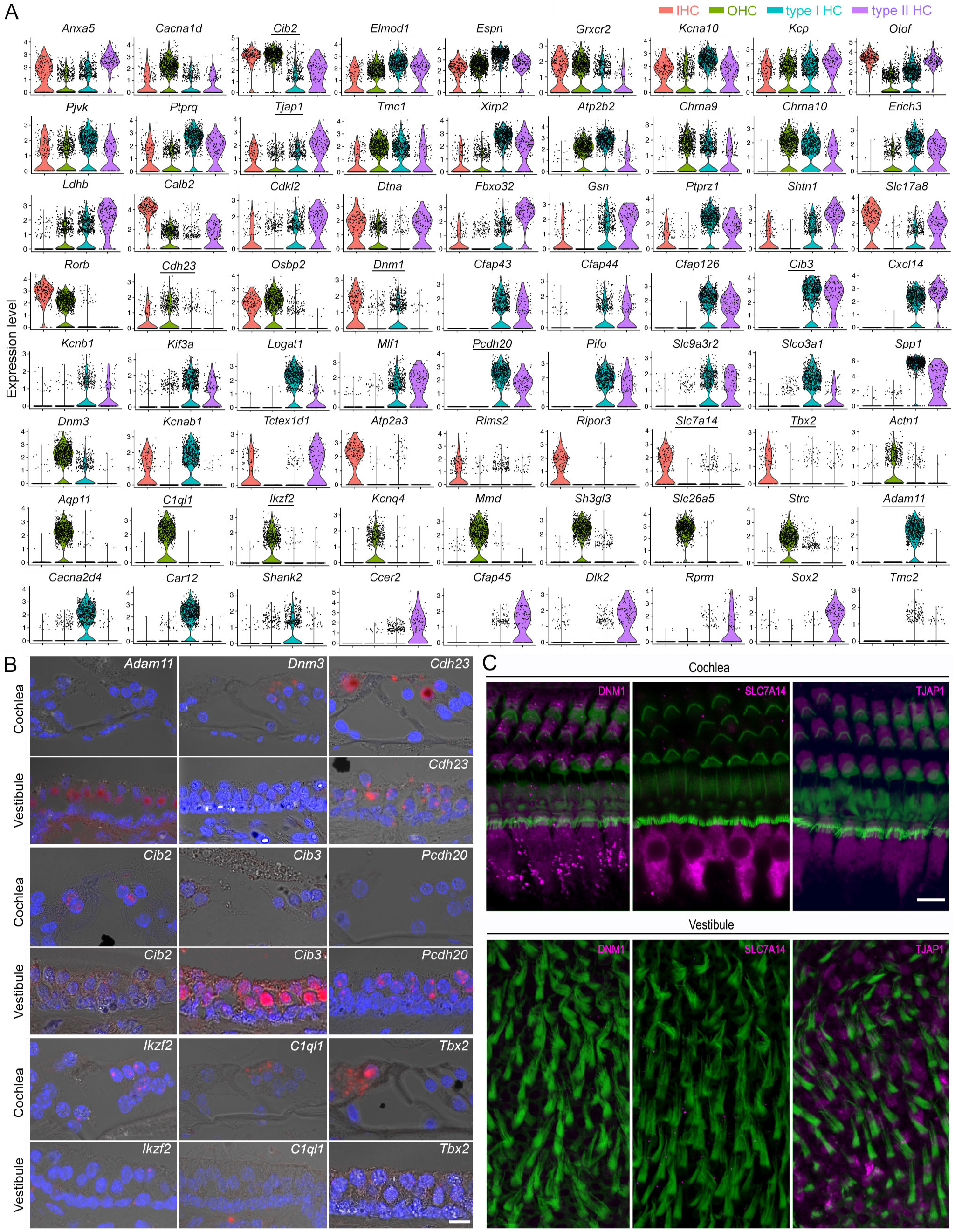
Shared and unique genes expressed in cochlear and vestibular HCs. **A**: Violin plots of the expression 72 genes in four different HC types. **B**: Validation of differential expressions of 9 genes (with underline in **A**) in cochlear and vestibular HCs in thin section. Bar: 10 μm for all images in B. **C**: Confocal images of expression of DNM1, SLC7A14, and TJAP1 in cochlear and vestibular HCs. Bar: 10 μm for all images in C.

Next, we focused our analysis on evaluating the genes associated with HC function. We compared the expression of 208 genes related to HC structure and function, including stereocilia and apparatus for mechanotransduction, ion channels, and synapses (Fig. 4A). Among the genes associated with stereocilia and mechanotransduction apparatus (Shin et al., 2013), *Calm1*, *Calm2*, *Eps8*, *Espn*, *Fbxo2*, *Dynll2*, *Ush1c*, *Ywhae*, *Tmc1*, *Tmie*, *Cdh23*, *Pcdh15*, *Ank1*, *Ank3*, and *Lhfpl5* were highly expressed in all four HC populations. Others were highly expressed in only one or two HC populations. *Atp2a3*, *Calb2*, and *Dpysl2* were highly expressed in IHCs, while *Lmo7*, *Ocm*, and *Strc* were highly expressed in OHCs. Moreover, expression of *Dpysl2*, *Cdh23*, and *Cib2* was higher in cochlear HCs than in vestibular HCs, while *Xirp2*, *Pls1*, *Slc9a3r2*, and *Tubb4b* were more highly expressed in vestibular HCs than in cochlear HCs. Some genes were uniquely expressed in either cochlear or vestibular HCs. For example, *Cib3* and *Tmc2* are expressed in vestibular HCs but not in cochlear HCs.

**Figure 4:**
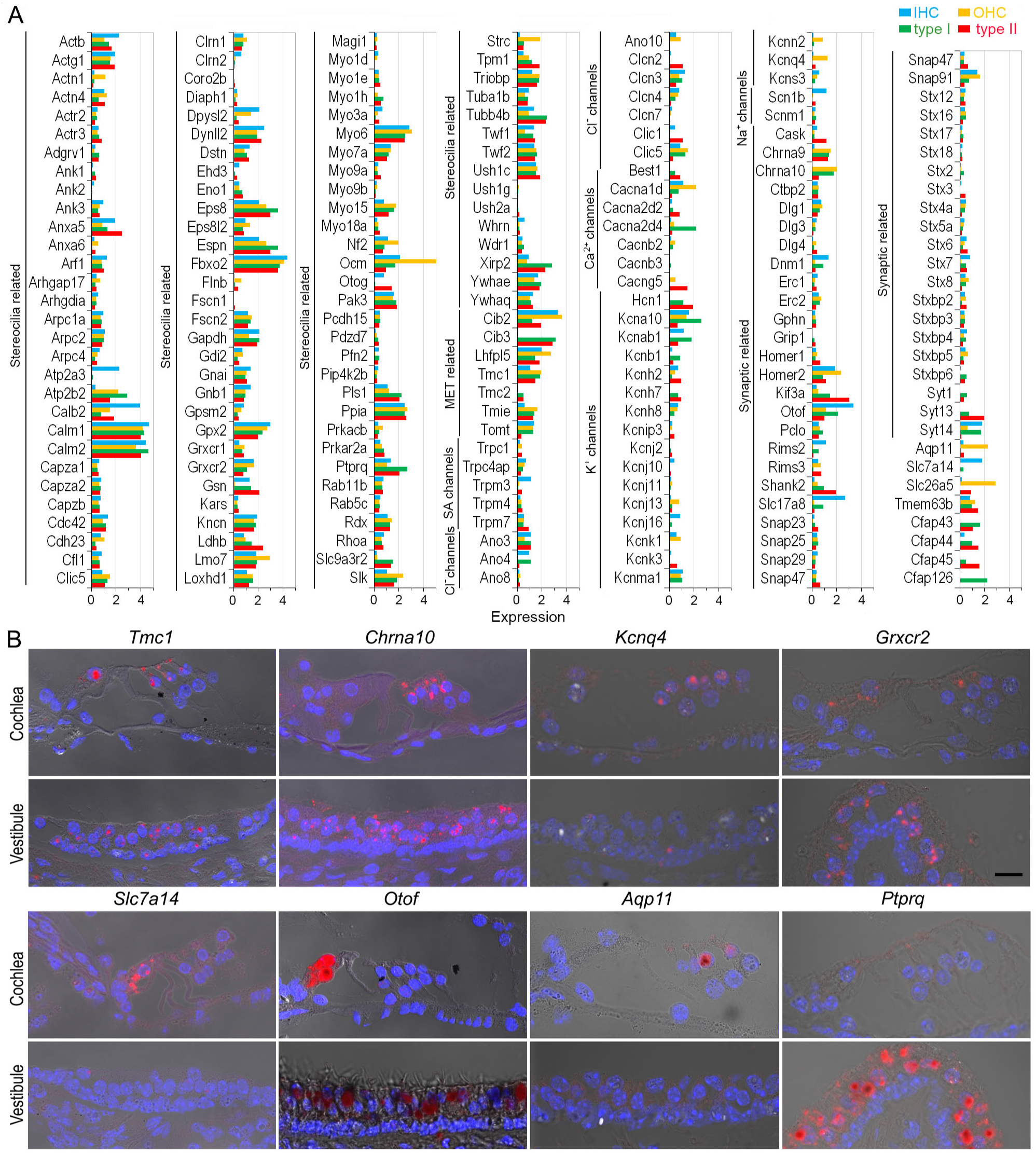
Expression of genes related to HC specialization. **A**: Heatmap showing expression of genes related to stereocilia bundles, mechanotransduction, ion channels and synaptic structure. **B**: Validation of gene expression using smFISH. Bar: 10 μm.

All HCs possess ion channels. Our analysis detected several genes related to stretch-activated ion channels, such as *Trpc1* and *Trpm4.* Genes for Cl^-^ and Na^+^ channels were expressed. For Ca^2+^ and K^+^ channels, *Cacna1d and Kcna10* were expressed in all four HC types with varying levels of expression. In cochlear HCs, *Cacna1d, Kcna10, Kcnab1, Kcnj16 and Kcnma1* were expressed in IHCs, whereas *Cacna1d, Kcna10, Kcnk1, Kcnma1, Kcnn2* and *Kcnq4* were expressed in OHCs. In the vestibular HCs, *Cacna2d4, Kcna10, Kcnab1,* and *Kcnma1* showed relatively high expression in type I HCs whereas type II HCs indicated a relatively high expression of *Cacna2d4, Cacng5, Kcna10, Kcnb1, Kcnh2,* and *Kcnh7*. *Best1, Clic, Hcn1,* and *Kcnh7* were only expressed in vestibular HCs.

Next, we examined the genes related to synapses (Fig. 4A). Our results indicated expression of *Ctpb2*, *Dlg1*, *Homer2*, *Otof*, *Pclo*, and *Snap91* in all HCs at varying levels. We observed a relative higher expression of *Dnm1*, *Otof*, *Rims2*, *Slc17a8*, *Snap91*, and *Stx7* in IHCs while *Dlg1*, *Dnm3*, *Snap9*, *Chrna9*, and *Chrna10* showed higher expression in OHCs. Some of the highly expressed genes in type I HCs include *Dnm1*, *Kif3a*, *Pclo*, *Shank2*, *Slc17a8*, *Syt13*, *Syt14*, *Chrna9*, and *Chrna10*, whereas type II HCs showed relatively higher expression of *Kif3a*, *Otof*, *Shank2*, *Stx7*, and *Syt13*. We employed smFISH to validate the expression of 8 additional genes across the four HC types. The expression patterns shown in Fig. 4B are consistent with our analysis (Fig. 4A).

### 4. Gene signatures of primary cilia in cochlear and vestibular HCs

A key morphological feature in the hair bundle of vestibular HCs is the presence of kinocilium, which has been regarded as a type of specialized primary cilia. Since our analysis revealed an enrichment of cilium-related GO terms (Fig. 2F) and axonemal genes such as *Cfap43, Cfap44, Cfap45, Cfap126, Kif3, Plf1,* and *Tubb4b* (Fig. 3A) in vestibular HCs, we sought to investigate the composition and molecular nature of the kinocilium (Fig. 5A). Proteomics-based approaches have contributed to the development of cilia-associated protein databases. We utilized several well-established databases, including CiliaCarta (Dam et al., 2019), the SYSCILIA gold standard (SCGSv2)(Vasquez et al., 2021), and CilioGenics (Pir et al., 2024) to compile a list of ∼1,000 cilia-related genes. We noted a significant overlap of these genes with our HC transcriptomic profiles (Fig. 5B). GO analysis of the overlapping genes revealed enrichment of cellular component and biological process terms primarily related to cilia organization, assembly, maintenance, intracellular transport, and microtubule dynamics particularly associated with motile cilia (Fig. S1). Additionally, molecular function analysis highlights associations with motor activity, dynein chain binding, and BBSome binding (Fig. S1). BBSome (Bardet-Biedl syndrome) is an octameric protein complex crucial for regulating transport in primary cilia (Tian et al., 2023).

**Figure 5.**
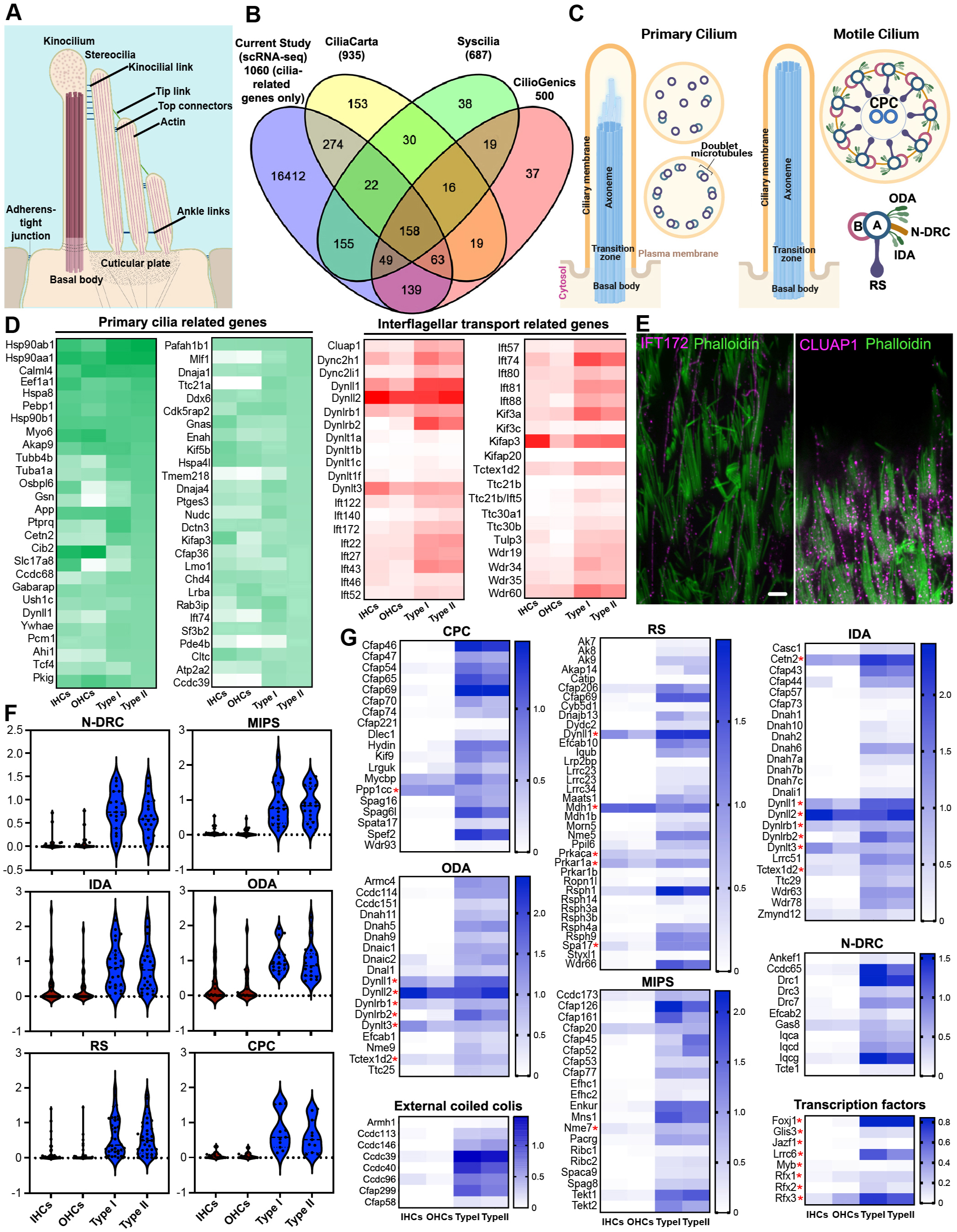
Cilia-related genes detected in cochlear and vestibular HCs. **A:** Schematic illustration of HC hair bundle (adapted from Schwander et al., 2010). **B**: Venn diagram of the number of genes in each database and the cilia-related genes detected in HC transcriptomes. **C**: Schematic illustration of primary and motile cilia, highlighting 9+0 or 9+2 arrangement of microtubules for primary and motile cilia, respectively: radial spokes (RS), central pair complex (CPC), nexin–dynein regulatory complex (N-DRC), microtubule inner proteins (MIPs), inner and outer dynein arms (IDA and ODA). **D**: Expression of top 50 cilia-related genes and genes related to IFT in the four types of HCs. **E**: Immunostaining of IFT172 and CLUAP1 expression in vestibular kinocilia. Bar: 5 µm. **F**: Violin plots showing aggregated expression of genes associated with 96-nm repeat. Expression value of these genes is based on Supplementary Table 1. **G**: Heatmaps of comparison of gene expressions related to motile-cilia machinery in cochlear and vestibular HCs. Red asterisks indicate the genes whose encoded proteins are expressed in both cilia and cytoplasm or are multifunctional.

The primary cilium is a sensory organelle that responds to and transmits external signals to the cell’s interior. Structurally, primary cilia are characterized by the presence of nine microtubule doublets (MTDs) encircling the shaft, which transition into a disorganized structure distally (Fig. 5C). We first examined the expression of primary cilia-related genes in vestibular and cochlear HCs. Among approximately 420 primary cilia-related genes/proteins (Dam et al., 2019; Pir et al., 2024; Vasquez et al., 2021), we detected the expression of 410 genes in at least one type of HC. Fig. 5D shows the top 50 abundantly expressed primary cilia-related genes in type II HCs compared to the other three HC types. Most of these genes were detected in all four HC types except a few genes, such as *Mlf1, Ttc21a,* and *Tmem218,* which were weakly or not expressed in cochlear HCs.

Primary cilia are enriched in receptors and effectors for key pathways, including GPCR, cAMP, Ca^2+^, RTK, TGF-β, MAPK, TOR, BMP, Wnt, Notch, and Rho signaling (Anvarian et al., 2019), localized to the ciliary shaft, transition zone, and BBSome (Hansen et al., 2025). We analyzed the expression of these genes in HCs and found higher expression levels of these pathway-related genes in vestibular HCs than in cochlear HCs (Fig. S2).

Intraflagellar transport (IFT) involves anterograde and retrograde transport of molecules along the axoneme of cilia, facilitating the transport of components between the ciliary base and tip (Ma et al., 2023). IFT is essential for the proper assembly and maintenance of both primary and motile cilia. Thus, we assessed the expression of IFT-associated genes in HCs. Most of the genes are enriched in vestibular HCs compared to cochlear HCs (Fig. 5D). Immunostaining confirmed the expression of IFT172 and CLUAP1 in the kinocilia of vestibular HCs (Fig. 5E).

### 5. Gene signatures of motile cilia in vestibular HCs

Curiously, our GO analysis returned many terms related to cilium motility. Motile cilia are highly conserved organelles across different organisms and tissues, although they exhibit organism- and tissue-specific adaptations (Leung et al., 2025). Recent advances using proteomics of isolated motile cilia from various ciliated tissues have enabled the profiling of genes associated with motile cilia. To explore whether the kinocilium possesses a molecular composition characteristic of motile cilia, we compared our HC transcriptomes with multi-tissue proteomics datasets derived from different organisms, including human, bovine, porcine, and murine, as well as diverse motile ciliary tissues such as sperm, oviduct, ventricle, and trachea (Leung et al., 2025) (Fig. S3). Our analysis revealed a significant overlap, particularly among axonemal components of motile cilia, with strong enrichment in vestibular HCs compared to cochlear HCs. The axoneme of motile cilia and flagella is a cylindrical structure harboring a canonical “9+2” arrangement, where nine doublet microtubules (DMTs) surround two microtubule singlets in the center (Fig. 5C). Altogether, the axoneme machinery consists of nine DMTs, two rows of inner and outer dynein arms (IDAs and ODAs), nexin-dynein regulatory complex (N-DRC), two singlet central pair complex (CPC), three radial spokes (RSs), microtubule inner proteins (MIPs), and external coiled-coil regions. The motility unit is arranged in a 96-nm repeat module along the CPC. Recent cryo-EM and cryo-electron tomography (cryo-ET) studies have provided a more comprehensive identification of this module repeat (Chen et al., 2023; Leung et al., 2025; Walton et al., 2023). To understand the molecular composition and function of kinocilia, we focused our analysis on the expression of genes associated with the structure of motile cilia. First, we assessed the expression of gene sets related to the motile cilium and each of its structural components. We noted a robust expression of motile cilia-related gene signatures in vestibular HCs, while cochlear HCs expressed little to none (Fig. 5F). Next, we further assessed the expression of the key genes related to motile cilia machinery (Fig. 5G), including the 96-nm module and CPC, based on the current known localization from biochemistry and proteomics.

We examined genes related to the 96-nm axonemal repeat of mammalian epithelial cilia. This structural unit contains proteins encoded by 128 genes (Gui et al., 2021). We found that 112 of these genes were expressed in adult vestibular HCs (Table S2). Genes encoding axonemal dynein (IDAs and ODAs), such as *Dnah5* and *Dnah6*, as well as RS components (*Wdr66, Cfap206, Cfap61, Iqub*), N-DRC components (*Drc1, Iqca*), and MIPs (*Cfap126, Wdr63*) were predominantly expressed in vestibular HCs with little to no expression in cochlear HCs (Fig. 5G). Axonemal CCDC39 and CCDC40, which form external coiled-coil regions, are the molecular rulers that organize the axonemal structure in the 96-nm repeating interactome and are required for the assembly of IDAs and N-DRC for ciliary motility (Becker-Heck et al., 2011; Brody et al., 2025; Merveille et al., 2011; Oda et al., 2014). Our results indicate a high expression of *Ccdc39* and *Ccdc40* in vestibular HCs, whereas little to no expression was observed in cochlear HCs. We should point out that unlike axonemal dynein proteins, which are uniquely required for cilia motility, the encoded proteins of several genes in Fig. 5G (marked by red asterisks) are also expressed in cytoplasm and/or are multifunctional.

Next, we examined expression of genes encoding transcription factors that are known key regulators of ciliome gene activation, including RFX and FOXJ1 transcription factor families. RFX controls genes in both motile and non-motile cilia, while FOXJ1 specifically governs motile cilia formation (Choksi et al., 2014). Our data show moderate expression of *Foxj1* in vestibular HCs and weak expression in cochlear HCs (Fig. 5G). Other genes that regulate motile cilia formation, including *Lrrc6*, were also expressed at high to moderate levels in vestibular HCs compared to cochlear HCs. We also found a strong enrichment of transcriptional targets associated with vestibular HCs, particularly those involved in motile cilia programming and maintenance. The expression of these transcription factors may reflect their importance in the maintenance of kinocilia in adult vestibular HCs. Furthermore, analysis of a published ATAC-seq dataset (Jen et al., 2019) from adult mouse vestibular tissue revealed increased chromatin accessibility in the promoter regions of genes associated with motile cilia machinery (Fig. S4), suggesting elevated transcriptional activity in vestibular HCs. These findings are consistent with our observations from scRNA-seq datasets.

To examine conservation of expression of these genes in vestibular HCs across species, we obtained the orthologs of axoneme-related genes in adult zebrafish inner ear HCs and human vestibular HCs using published datasets (Barta et al., 2018; Wang et al., 2024). Fig. 6A shows the expression of these genes in adult mouse, zebrafish, and human vestibular HCs. While the expression level varies, most of these genes are expressed across species. The exceptions are *Dnah3* and *Dnah12*, which are expressed in zebrafish HCs but not in mammalian vestibular HCs. We conducted immunostaining and high-resolution confocal imaging to validate the expression of key motile cilia markers in the kinocilia. FOXJ1 is expressed in the nucleus of vestibular HCs, while CCDC39, CCDC40, TEKT1, DNAH5, and DNAH6 are expressed in kinocilia (Fig. 6B). Collectively, our findings provide evidence corroborating the presence of motile cilium machinery in the kinocilia of vestibular HCs.

**Figure 6:**
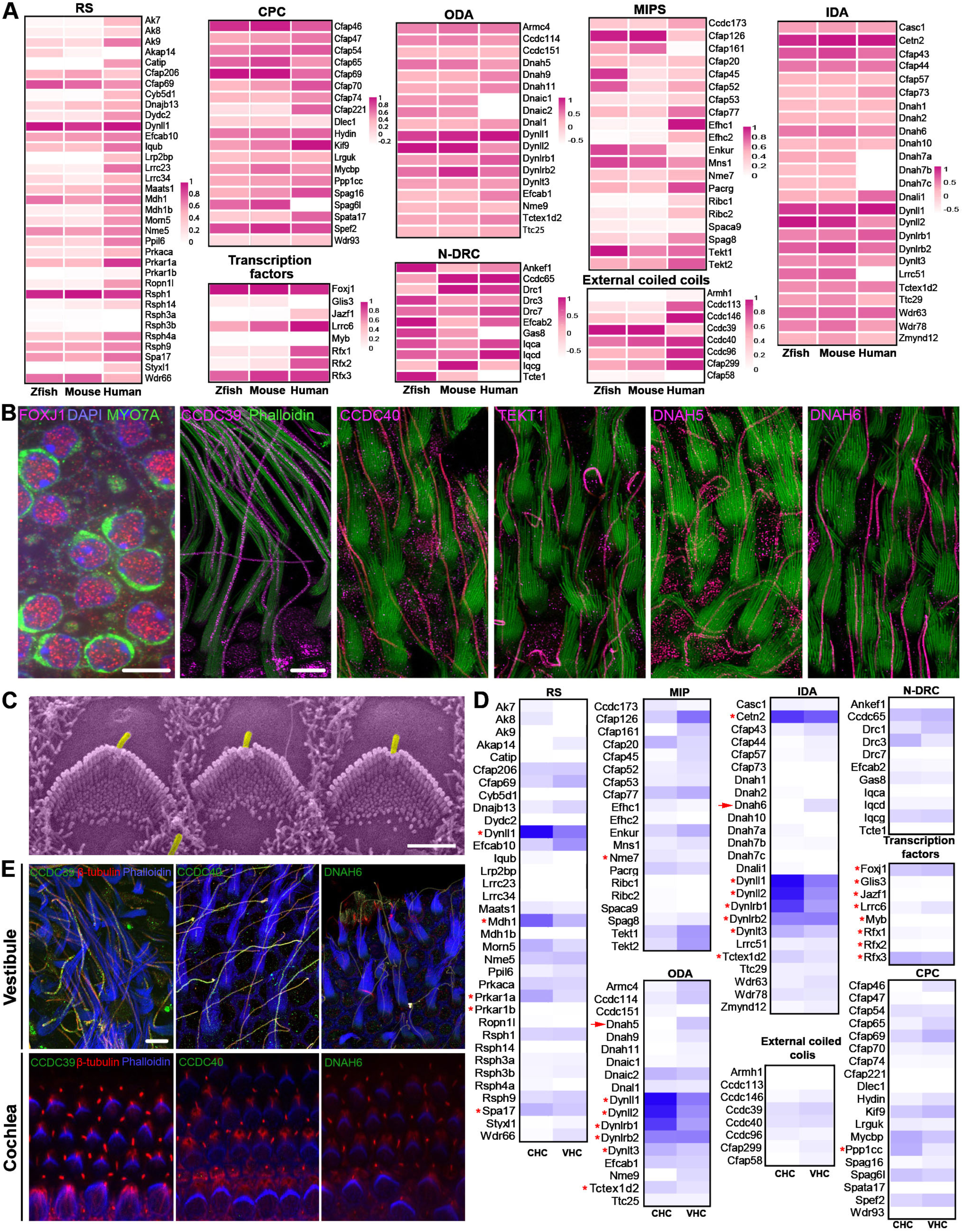
Expression of motile cilia-related genes/proteins in the vestibular HCs. **A:** Expression of motile cilia-related genes in zebrafish, mouse, and human vestibular HCs. Expression values of these genes are based on Supplementary Table 1 (mouse), Data Citation 4 (zebrafish, Barta et al., 2018), and Wang et al., 2024 (human). Mouse gene nomenclature is used in heatmaps. **B:** Confocal images of the expression of key motile cilia-related proteins. Scale bars represent 5 µm. **C**: SEM micrograph of hair bundles of OHCs from P2 cochlea. Kinocilia (in magenta) are still present at this age. Bar: 2.5 μm. **D:** Comparison of expression of motile cilia-related genes between P2 cochlear and vestibular HCs. Gene expression values are based on HC transcriptomic dataset by Burns et al., 2015. Red asterisks mark the genes whose encoded proteins are expressed in both cilia and cytoplasm or multifunctional. Red arrows indicate *Dnah5* and *Dnah6*, which were not detected in P2 cochlear HCs. **E:** Confocal images of expression of CCDC39, CCDC40 and DNAH6 in P2 vestibular and cochlear HCs. CCDC39, CCDC40, and DNAH6 were not expressed in cochlear HCs at P2. Bar: 5 μm.

Since nascent cochlear HCs possess kinocilia (Fig. 6C), we used published P2 cochlear and vestibular HC transcriptomes (Burns et al., 2015; McInturff et al., 2018) to investigate whether neonatal cochlear HCs express motile cilium-related genes. Assessment of expression of genes related to motile cilia machinery revealed a less drastic difference between neonatal cochlear and vestibular HCs than that between adult cochlear and vestibular HCs (Fig. 6D). We note that proteins of some shared genes are expressed in both cilia and cytosol (marked by red asterisks in Fig. 6D). However, some key motility-associated genes such as *Dnah6* and *Dnah5* (marked by red arrows in Fig. 6D) were not detected in the P2 cochlear HCs. These axonemal dynein heavy chains are ATPase-driven force-generating motors that produce the ciliary power stroke in concert with other axonemal components. Immunostaining confirmed the lack of expression of CCDC39, CCDC40, and DNAH6 in cochlear HCs at P2 (Fig. 6E). In contrast, these key proteins were expressed in kinocilia of vestibular HCs (Fig. 6E). Lack of expression of *Dnah5* and *Dnah6* and the molecular rulers CCDC39 and CCDC40 suggests that the kinocilium of neonatal cochlear HCs does not possess signatures of motile cilia. Therefore, our analysis indicates that the molecular composition of kinocilia is different between neonatal cochlear and vestibular HCs.

### 6. Vestibular kinocilia exhibit hybrid morphological features of primary and motile cilia

TEM studies have characterized the ultrastructure of kinocilia across species, revealing evidence of complex and regionally specialized organization (Kikuchi et al., 1989; Nagel et al., 2014; O’Donnell and Zheng, 2022). We used TEM to examine the ultrastructure of kinocilia from bullfrog crista HCs. TEM images, including longitudinal sections of frog vestibular hair bundles (Fig. 7A) highlight a distinct zonal architecture along the kinocilium axis. At the distal tip, a prominent kinociliary bulb is observed, while the base anchors the axoneme within the cuticular plate. Two pairs of microtubule doublets span the full length of the kinocilium in the longitudinal section. The central pair of singlet microtubules, typical of motile cilia, is maintained throughout most of the shaft but disappears in the distal and transitional zones. This configuration results in a dynamic shift from a canonical 9+2 arrangement centrally to a 9+0 pattern at both the base and tip. These observations underscore the heterogeneous and hybrid nature of vestibular kinocilia, integrating structural features of both primary and motile cilia.

**Figure 7:**
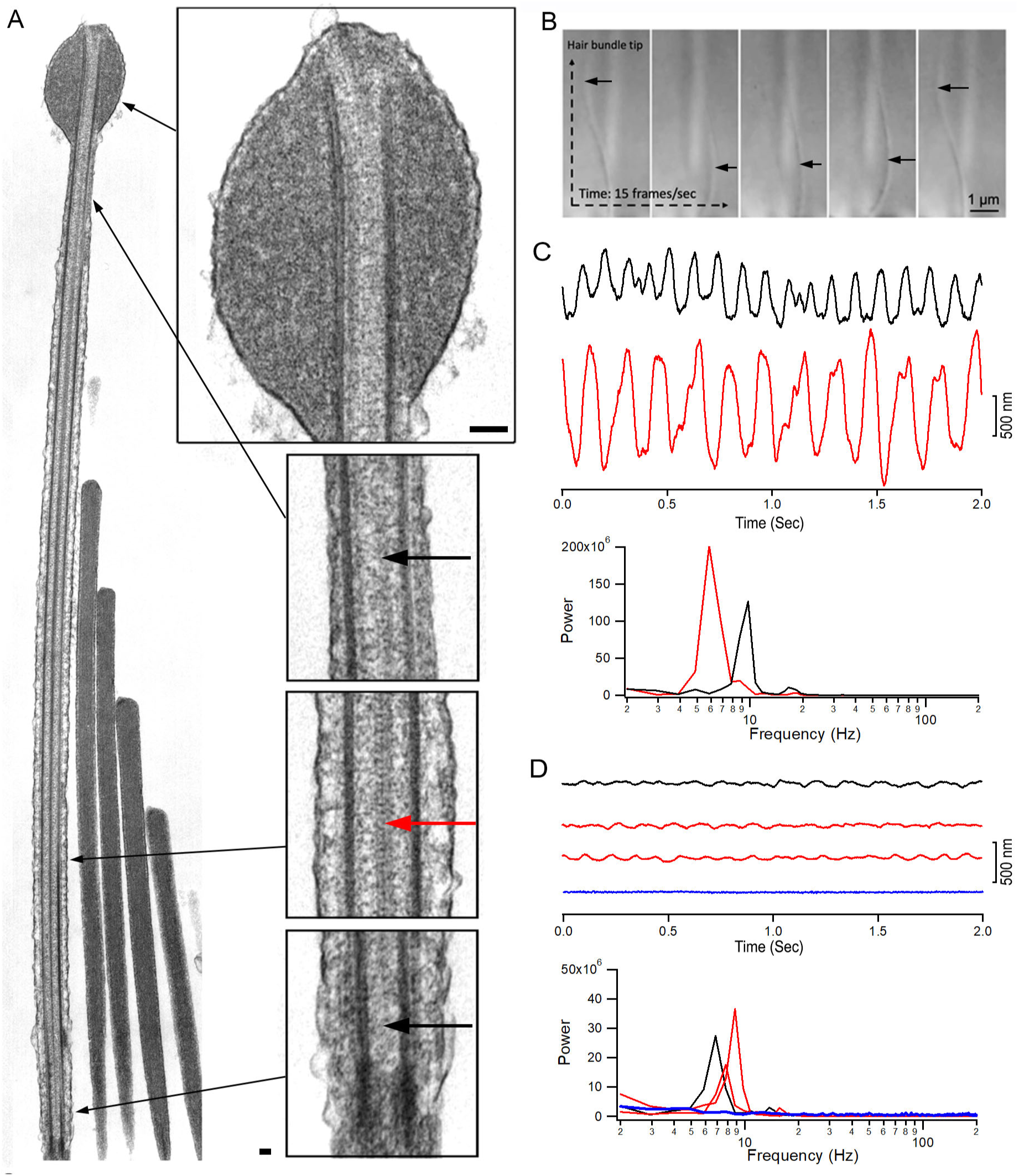
Kinocilia morphology and motility. **A**: TEM images of stereocilia and kinocilium from bullfrog crista HCs. Different regions of the kinocilium in higher magnification are also shown. Long black arrows indicate where the magnified images were taken. Bars: 250 nm. Red arrow indicates two central microtubule singlets. Short black arrows mark absence of central microtubule singlets in the distal regions near the tip of kinocilium and transition zone. **B**: Images captured from *in vitro* live imaging of kinocilium and bundle motion of a bullfrog crista HC. The images were captured at a speed of 15 frames per second. Black arrows indicate kinocilium. **C**: Representative waveforms of spontaneous cilia motion from middle ear tissue. The FFT analysis of cilia motion is also shown. **D:** Three representative waveforms of spontaneous motion of hair bundles. The response waveform in blue was taken from a hair bundle with no spontaneous motility. FFT analysis of bundle motion is shown. Response waveforms and spectra are color-coded and -matched.

### 7. Evidence that vestibular kinocilia exhibit motility

Vestibular kinocilia are traditionally regarded as non-motile, lacking the rhythmic beating characteristic of respiratory cilia. Kinocilia are not only tightly connected to stereocilia but are also embedded in the overlaying gelatinous membrane (Eatock and Songer, 2011; Leibovici et al., 2005; Li et al., 2008), an extracellular matrix required for physiological mechanotransduction. Using acute preparations of bullfrog semicircular canal sensory epithelia (cristae), we observed robust spontaneous kinociliary motility in some HCs (Fig. 7B; Videos S1 and S2). This motility exhibited the characteristic beating pattern of flagella and cilia and occurred at a frequency of approximately 5 - 10 Hz at room temperature. The observed displacements were sufficiently large to induce deflection of the entire hair bundle, indicating it can generate forces to influence hair bundle dynamics. Interestingly, this phenomenon was detected in only ∼1 - 5% of crista HCs, likely due to variable preservation of HC and kinociliary integrity *in vitro* in acutely dissected tissue. These findings suggest that vestibular kinocilia are capable of active movement and challenge the strict classification of these structures as non-motile even though motility is not experimentally observed in most HCs in the excised vestibular sensory epithelium.

Next, we explored whether the kinocilia of vestibular HCs from adult mice are motile by utilizing the photodiode technique to detect bundle motion (Jia and He, 2005). This technique can detect motion in the 10-nm range for synchronized signals after averaging. Since spontaneous motions are not synchronized and cannot be averaged to improve signal-to-noise ratio, we captured the responses in the time domain and averaged them in the frequency domain after a fast Fourier transform. This allows us to detect unsynchronized cilia motion even if the signal is close to the noise level at the time domain. We first measured spontaneous cilia movement from the epithelial lining of the Eustachian tube, which is an extension of airway epithelia, as cilia in the respiratory tract are a typical example of motile cilia. Fig. 7C shows two representative waveforms of airway cilia beat with a magnitude between 700 and 1500 nm. The two responses shown in Fig. 7C have main frequency components at 6 to 9 Hz at room temperature. Next, we measured the movement of the top segment of the hair bundle from mouse crista ampullaris. Since the kinocilium is tightly attached to the stereocilia bundle, we measured the whole bundle’s motion due to the difficulty of taking measurements from a single kinocilium. The waveform (black trace in Fig. 7D) was obtained from crista HCs bathed in perilymph-like solution (L-15 medium) with 2 mM of Ca^2+^. Like the beat frequency of airway cilia, kinocilia also moved at the frequency of ∼7 Hz (Fig. 7D). To rule out the possibility that the bundle motion is driven by the mechanotransduction-related activity (Benser et al., 1996; Martin et al., 2003), we treated the crista ampullaris in Ca^2+^-free medium with EGTA for 2 minutes to break the tip-link (Assad et al., 1991; Jia et al., 2007; Kachar et al., 2000; Ricci et al., 2003). Spontaneous motion was still detected (two red traces in Fig. 7D), suggesting that the motion is independent of the transduction channel activity. We measured spontaneous bundle motions from 52 crista HCs from six mice. Spontaneous motion was only detected in eight HCs. An example of a lack of response is shown in Fig. 7D (blue trace). Although we were unable to determine which of the HC subtypes were exhibiting kinocilia motility, it is conceivable that both type I and type II HCs have this capability since they both express the genes related to motile cilia. We note the kinocilia motion of mouse crista HCs was substantially smaller than that of airway cilia (Fig. 7D) and bullfrog crista HCs.

### 8. Predicted model of the 96-nm modular repeat in adult vestibular kinocilia

Since kinocilia motility in mouse vestibular HCs is substantially smaller than the motility of airway cilia, we investigated the structural basis underlying comparatively reduced motility of kinocilia. This diminished motility may result from differences in the molecular composition and organization of the axonemal machinery, particularly the 96-nm modular repeat that houses key dynein motors and regulatory complexes. Recent advances in structure prediction powered by artificial intelligence and cryo-electron microscopy (cryo-EM) have facilitated the generation of highly conserved atomic models of the 96-nm axonemal repeat from human respiratory cilia and bovine sperm flagella (Chen et al., 2023; Leung et al., 2025; Walton et al., 2023). We applied our axonemal gene dataset to these atomic models to predict the molecular architecture of the 96-nm repeat in vestibular kinocilia. By mapping the expression of known axonemal components onto the structural frameworks derived from human respiratory (PDB: 8J07) (Walton et al., 2023) and bovine sperm (PDB: 9FQR)(Leung et al., 2025) axonemes, we generated two composite models that reflect the unique molecular composition of the vestibular kinocilium. These predicted structures are shown in Figs. 8A and 8B. 3D videos of the predicted models are provided in Video S3 and Video S4.

**Figure 8:**
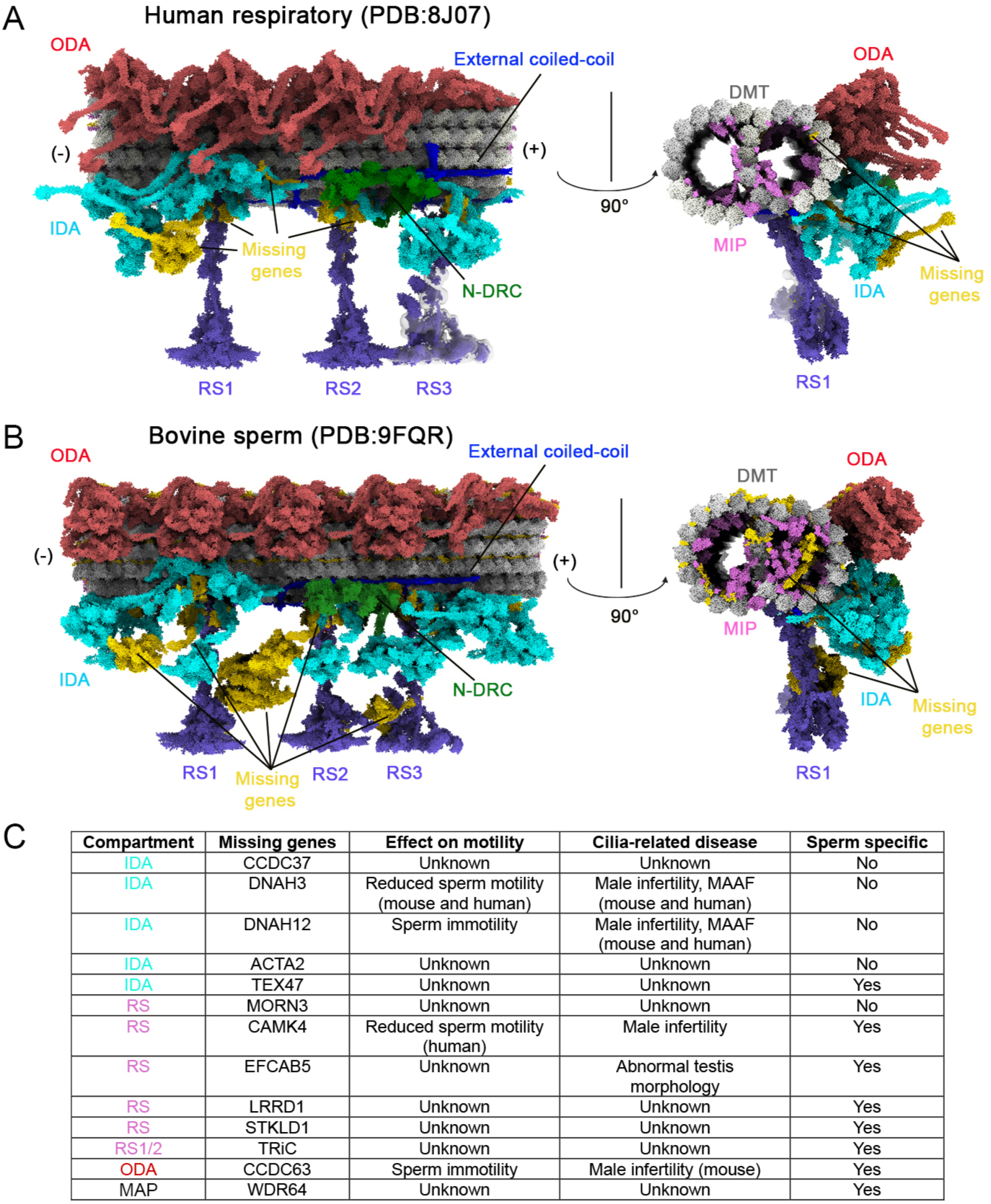
Predicted models of the molecular architecture of 96 nm axonemal repeat of vestibular kinocilia. **A-B**: Longitudinal and cross-sectional views of the doublet microtubule and associated structure in 96-nm repeat, derived from combining cryo-EM data and single-cell transcriptomic analysis from human respiratory (A) and bovine sperm flagella (B). Key axonemal motile-machinery components are color-coded: ODA (Indian red), IDA (cyan), N-DRC (green), MIPs (orchid), RS (purple), and external coiled-coils (blue). Radial spoke 3 (RS3) has not been resolved to atomic resolution, but its shorter form (RS3s) is depicted. Doublet microtubules (DMT) are represented in gray. Regions highlighted in gold indicate the absence of corresponding transcripts in our mouse transcriptomic data. **C**: Genes which are not detected in mouse and human vestibular HC transcriptomes and related to motility-relevant compartments are listed in the table. The roles of these genes in the 96-nm repeat module and cilia motility and ciliopathy are also included.

We chose the human respiratory 96-nm axonemal structure as our reference because it best reflects vestibular kinocilia and provides a more comprehensive representation of axonemal components than the sperm model, which shows more missing elements (highlighted in gold in Figs. 8A,B). Using a curated reference gene list derived from the human respiratory 96-nm axonemal structure, we mapped vestibular HC gene expression and identified transcripts for all 18 ODA genes and their docking complex components, all 11 N-DRC genes, and all 7 MAPs, along with 36 of 37 RS genes and 19 of 23 IDA-related genes. Additionally, 20 of the 31 genes encoding MIPs, which are known to stabilize the microtubule doublets, were also expressed (Fig. 5G). In contrast, vestibular HCs lacked the expression of several sperm-specific genes from distinct compartments of 96-nm repeat (Leung et al., 2025), including TRiC chaperonin subunits (Brown et al., 2025; X. Meng et al., 2024), CAMK4, EFACB5, LRRD1, STKLD1, CCDC63, WDR64, and several MIPs (Fig. S5).

Based on our models, the absence of *Dnah3* and *Dnah12*, along with *Acta2*, is predicted to result in the loss of two of the six single-headed IDA components (highlighted in gold). *Cfap100*, which forms the modifier of inner arms (MIA) complex with *Cfap73* and contributes to tethering the double-headed IDAf (inner dynein arm f), is missing in the model, whereas *Cfap73* and the remaining IDAf components are present. Notably, IDAf has multiple attachment points, and MIA is not essential for docking IDAf to the DMTs (Yamamoto et al., 2013). Overall, 4 of the 23 IDA-related genes were not expressed in vestibular HCs; nonetheless, we predict that the IDA structure is reduced but not entirely absent, as DNAH6 remains localized in vestibular kinocilia (Fig. 5B). In addition to the missing IDA components, we identified 11 unexpressed genes associated with MIPs, whose absence is predicted to result in reduced MIP density in the models (highlighted in orchid and gold in cross-sectional views in Figs. 8A, and B). Unlike other axonemal structures, MIPs exhibit greater variability across species, which may account for their lineage-specific absence in vestibular kinocilia (Fig. 8)(Andersen et al., 2024; Tai et al., 2023; Xia et al., 2025). The missing genes in our HC datasets and their roles in the axonemal complex, cilia motility and ciliopathy are listed in Figs. 8C and S5. Based on our predicted models, we speculate that the absence of *Dnah3* and *Dnah12* plays a major role in limiting kinocilia motility in mouse vestibular HCs, contributing to the smaller movements compared to motile cilia.

## DISCUSSION

This is the first study to compare transcriptomes among four types of HCs from the adult mouse inner ear and to characterize the molecular composition of kinocilia. We found that the transcriptomic similarity between type I and type II HCs is greater than that between IHCs and OHCs, indicating greater homogeneity among vestibular HC subtypes compared to cochlear HC subtypes. We identified several new genes and proteins that can be used as markers for vestibular HCs, especially those related to kinocilia. We observed notable differences in gene signatures related to HC unique structure and function, which may underlie distinct biological properties of mechanotransduction, membrane conductance, and synaptic transmission seen among the four different HC types. Differential expressions also explain why loss of function of a gene such as *Tmc1*, *Cib2*, or *Cib3* leads to differential auditory and vestibular phenotypes in mouse models and humans. Our dataset is expected to serve not only as a valuable resource for unraveling the molecular mechanisms of the biological properties of HCs but also for assisting the auditory and vestibular research community in identifying and exploring the functions of disease-related genes. Since biological processes enriched in vestibular HCs are related to cilia and cilia motility, we focused our analyses on kinocilia. Although the kinocilium has long been considered a primary cilium, its molecular composition and structural organization remain largely unexplored. Our study suggests that kinocilia serve dual roles as both primary and motile cilia. The primary cilium is a major hub in receiving and transmitting signals from the environment. A recent study has demonstrated the expression of proteins related to transduction pathways and receptors in primary cilia using spatial proteomics (Hansen et al., 2024). In line with this, we found that both vestibular and cochlear HCs express genes encoding components of signal transduction pathways and receptors. While the precise localization of these proteins within HC kinocilia remains to be validated, our analysis reveals a shared expression of diverse primary cilium signaling genes across HC subtypes. This suggests a conserved, and potentially specialized, role for kinocilia in cellular signaling and ciliary function.

Although adult cochlear HCs lack kinocilia, we still observed expression of numerous cilia-related genes in in these cells (Figs. 5 and 6). While some of these genes may be vestigial, many are associated with primary cilia structures, including the basal body and IFT machinery. Notably, the basal body persists in adult cochlear HCs despite the developmental disappearance of the kinocilium. Previous studies using cilia proteomics have shown that many cilia-related proteins are expressed in cytosol, including proteins related to signal transduction, microtubule cytoskeleton, actin cytoskeleton, vesicle transport, metabolism, protein folding, translation, nuclear transport, ubiquitination, RNA binding, mitochondria, and transcriptional regulation (Dam et al., 2019; Pir et al., 2024). Therefore, it is not unexpected that adult cochlear HCs continue to express genes associated with ciliary functions.

A TEM study of mouse vestibular kinocilia (O’Donnell and Zheng, 2022) showed the ‘9+2’ microtubule arrangement characteristic of motile cilia. This contrasts with an earlier study of guinea pig vestibular kinocilia, which reported the absence of central singlet microtubules and IDAs, while ODAs and RSs were present (Kikuchi et al., 1989). In our study, TEM revealed a complex and regionally specialized organization of the kinocilium in bullfrog vestibular HCs. The axoneme is anchored within the cuticular plate at the kinocilium base. While the central pair of singlet microtubules is preserved along most of the shaft, it is absent in the distal and transitional zones—indicating a transition from the canonical 9+2 microtubule arrangement to a 9+0 configuration at both the base and tip. In most motile cilia, the central pair does not originate directly from the basal body; instead, it begins a short distance above the transition zone, a feature that illustrates variation across systems (Lechtreck et al., 2013). The central pair can also show variation in its spatial extent: for example, in mammalian sperm axonemes, it can terminate before reaching the distal end of the axoneme (Fawcett and Ito, 1965). In addition, the central pair orientation differs across organisms: in metazoans and *Trypanosoma,* the central pair is fixed relative to the outer doublets, whereas in *Chlamydomonas* and ciliates it twists within the axoneme (Lechtreck et al., 2013). Such structural variation has been observed in various motile cilia and flagella and is therefore not unique to vestibular kinocilia. However, a more distinctive feature of kinocilium morphology is the organization at the distal tip, where a prominent distal head is present—resembling tip structures recently identified in human islet cell cilia (Polino et al., 2023).

This distal-most region is known to harbor specialized proteins (Legal et al., 2023). In multi-ciliated cells, CCDC33 and CCDC78 are found at the very end of the cilium and help organize other proteins like SPEF1, CEP104, and EB3/MAPRE3 (Hong et al., 2025; Legal et al., 2024, 2023). We observed expression of all genes encoding these proteins, except for *Ccdc78*. Although we did not study the tip of the kinocilium, its bulb shape suggests it may also contain specialized proteins. In bullfrog HCs, the kinocilial bulb binds to the overlaying otoconial membrane (Jaeger et al., 1994; Kachar et al., 1990), and shows strong labeling for β-tubulin and cadherin 23 (Jaeger et al., 1994; Kachar et al., 1990; Lagziel et al., 2005), and a recent study showed that *saxo2* overexpression in zebrafish results in bulbed kinocilia, with *saxo2* protein accumulation at the distal tip (Erickson et al., 2023). These bulbed tips may reflect specialized regulation of ciliary cap proteins (Legal et al., 2023) that help organize or stabilize the plus ends of axonemal microtubules—an area that remains to be explored in kinocilia.

The most important and novel finding of our study is that adult mouse vestibular HCs express genes related to the 96-nm axonemal repeat complex, a hallmark structural feature of motile cilia. Notably, orthologs of these genes are also expressed in zebrafish and human vestibular HCs, highlighting the evolutionary conservation of this molecular complex across vertebrate species. Furthermore, we observed robust spontaneous kinociliary motility in bullfrog crista HCs, as well as subtle spontaneous bundle movements in mouse crista HCs. Our findings indicate that this motion is independent of the mechanotransduction apparatus, as the kinocilium itself exhibited active, flagellar-like movement in bullfrog crista HCs (Video S1 and Video S2). In mouse crista HCs, spontaneous motion was still present after breaking the tip links. Early studies reported observation of spontaneous flagella-like rhythmic beating of kinocilia in vestibular HCs in frogs and eels (Flock et al., 1977; Rüsch and Thurm, 1990), as well as in zebrafish HCs in the early otic vesicle (Stooke-Vaughan et al., 2012; Wu et al., 2011). According to Rüsch and Thurm, spontaneous kinociliary motility was observed only under conditions of tissue deterioration. They therefore interpreted kinocilia beating as a sign of cellular decline rather than a physiological feature. We speculate that such deterioration may have disrupted kinocilial links, effectively unloading the kinocilium and permitting freer movement. Nonetheless, the observation of spontaneous kinocilia beating—regardless of tissue condition—supports the conclusion that they are motile cilia.

We observed kinociliary beating in only a subset of the cells. While we cannot exclude the possibility that indeed only some kinocilia are inherently motile, or that stressful conditions may activate a latent motility in vestibular HCs, there are several possible reasons why such motility has not been consistently observed in most vestibular HCs – both in our study and in previous investigations. For example, a reduction in intracellular ATP levels in *in vitro* preparations may play a significant role, as ciliary motility is ATP-dependent. Rapid depolarization of HCs *in vitro* is an indication of reduced availability of intracellular ATP, as Na^+^/K^+^-ATPase pump is critical for maintaining resting membrane potential (He and Dallos, 1999; Silver and Erecińska, 1997). Additionally, acute tissue dissection may disrupt kinocilia, which are normally straight and tethered to the extracellular matrix. *In vitro*, many appeared bent, potentially compromising their structural integrity and the metabolic conditions required for motility.

Although we observed kinocilium-driven bundle movement in adult mouse vestibular HCs, its magnitude was at least an order of magnitude lower than that of other motile cilia. Our single-cell transcriptomic analysis showed the absence of 16 genes associated with the 96-nm axonemal repeat of human respiratory cilia, including *Pierce1* and *Pierce2*. These two MIPs-encoding genes have been shown to regulate motile cilia function and left–right asymmetry in mouse models (Gui et al., 2021; Sung et al., 2016). *Pierce1* knockout results in pronounced defects in ciliary motility and dynein arm docking, while *Pierce2* loss has a milder effect and largely preserves overall ciliary ultrastructure and beating (Gui et al., 2021; Sung et al., 2016). Although the contribution of these genes to kinociliary function remains uncertain, the absence of both may contribute to reduced microtubule stability or motor organization, especially when combined with additional losses in key motor components. However, MIPs are the most heterogeneous components across different types of cilia, such as sperm and airway cilia in different species (Leung et al., 2025; Tai et al., 2023). Thus, it remains unclear whether the absence of these two genes reduces kinociliary motility in mouse vestibular HCs.

The lack of expression of two genes associated with IDAs (*Dnah12*, *Dnah3*) may lead to the loss of specific single-headed IDA components as suggested by our structural models (Fig. 8). Mutations of *DNAH3* and *DNAH12* are linked to male infertility and dynein dysfunction in humans (Meng et al., 2024; Yang et al., 2024). Mutations of DNAH12 cause male infertility by impairing DNAH1 and DNALI1 recruitment. However, it does not affect the tracheal tract and oviductal cilia organization (Yang et al., 2024). For the other two IDA-related genes (*Acta2, Cfap100*/*Ccdc37*) and one RS-related gene (*Morn3*), no ciliopathy has been linked to mutations of these three genes so far. We speculate that the lack of expression of DNAH3 and DNAH12 may be a key factor limiting magnitude of kinocilia motility in mammalian vestibular HCs compared to respiratory cilia or kinocilia of bullfrog vestibular HCs.

We detected the expression of a few sperm-specific MIPs including *SAXO4*, *Tektin 3*, and *Tektin 4* in vestibular HCs. While the kinocilium shares a broader molecular profile with epithelial motile cilia, the presence of these distinct sperm-specific MIPs, which are absent from multi-ciliated epithelial tissues, suggests kinocilia may possess unique structural specializations adapted to their exceptional length (∼60 – 70 μm in length in mouse crista HCs) and sensory function, highlighting unique identity of kinocilia.

HCs employ positive local feedback to amplify inputs to their mechanosensitive hair bundles. This amplification helps overcome mechanical impedances and fine-tune sensory stimuli. In mammals, the remarkable sensitivity of the auditory system is largely attributed to the fast somatic motility of OHCs in the cochlea (Brownell et al., 1985; Dallos et al., 2008; Kachar et al., 1986; Liberman et al., 2002; Zheng et al., 2000). In other receptor organs, HCs may effect amplification by the Ca^2+^-dependent activity of myosin or transduction channels in the stereocilia (Fettiplace, 2017; Hudspeth, 1997). Mechanotransduction-mediated active hair bundle movements have been reported in turtle and frog HCs (Benser et al., 1996; Crawford and Fettiplace, 1985; Denk and Webb, 1992; Howard and Hudspeth, 1987; Martin et al., 2003). The functional significance of kinociliary beating remains to be elucidated; however, the kinocilium may serve as an active, force-generating component of the hair bundle. Because the kinocilium is connected to the tallest stereocilia via kinocilial links, we speculate that kinociliary motility may dynamically modulate the mechanical properties of the hair bundle or influence tip-link tension to prime transduction channels. Kinociliary beating is sufficient to drive stereocilia bundle movement (Videos S1 and S2). Even when constrained by the overlying otolithic membrane or cupula, changes in kinociliary stiffness could still affect the bundle’s mechanical dynamics. Importantly, such autonomous rhythms are unlikely to disrupt temporally accurate encoding of head motion, as spontaneous bundle movements driven by mechanotransduction have also been observed in bullfrog saccular HCs (Benser et al., 1996; Martin et al., 2003). Moreover, auditory and vestibular afferent neurons also generate spontaneous action potentials in both developing and mature animals. Although we did not examine when spontaneous kinocilia beat emerges during development, our analysis showed that key motile cilia signature genes and proteins are expressed at P2, suggesting that spontaneous kinocilia beat may already be present at this age. Such activity may help refine HC maturation and neural connections and prime the vestibular central pathway for its later function.

In summary, this study demonstrates that the kinocilium of vestibular HCs is a unique hybrid cilium, exhibiting near-complete molecular features of both primary and motile cilia. While it shares structural and molecular similarities with motile cilia and sperm flagella, it also possesses distinct architectural and functional characteristics. Future investigations employing kinocilium-specific proteomics, cryo-ET, and single-particle analysis will be critical for fully characterizing kinocilium molecular composition and organization. Although kinocilia motility was observed in bullfrog and mouse vestibular HCs in the present study, the functional significance of kinocilia motility remains to be elucidated.

## MATERIALS AND METHODS

Male and female CBA/J mice were purchased from the Jackson Laboratory (Stock #:000656) and reared in Animal Care Facility of Creighton University and NIDCD. American bullfrogs (*Rana catesbiana*) were purchased from Carolina Biological Supply Co. The animal usage and care were approved by the Institutional Animal Care and Use Committees of Creighton University (Protocol #1046.3) and NIDCD (NIDCD ACUC Protocol #1215).

### 1. Cell dissociation, cDNA libraries preparation and RNA-Sequencing

Male and female CBA/J mice aged 10 weeks were used for scRNA-seq. Cochlear and vestibular end organs (utricle, saccule, and crista) were dissected from the inner ear and placed in Petri dishes containing cold L-15 medium (Gibco; #11320033). After the cochlear and vestibular sensory epithelia and neurons were dissected out, they were transferred into two individual 1.5 ml tubes for enzymatic digestion (Collagenase IV from Sigma, concentration: 1mg/ml collagenase) in L-15 medium. After 10 minutes of incubation at room temperature, the enzymatic solution was removed and replaced with 400 µl L-15 media containing 10% fetal bovine serum. The tissues in two tubes were mechanically triturated by 200 µL Eppendorf pipette tips. After that, the suspension containing cochlear and vestibular cells was then passed through 40 µm strainers for filtration and pelleted at 300 g for 5 mins. After removing extra media, cells were then reconstituted in the 50 ul L-15 with 10% fetal bovine serum media and used for cDNA library preparation. Seven mice were used for each biological replicate. Six biological replicates for cochlear sensory epithelium and four biological replicates for vestibular sensory epithelia were prepared for scRNA-seq.

The emulsion droplets were constructed using a 10x Genomics Controller device following the manufacturer’s instruction manual. cDNA libraries were constructed using the 10x Genomics Chromium Single Cell 3’ Reagent Kits V3.1. High Sensitivity DNA Kits (Agilent Technologies) were used to perform quality control for each library in an Agilent 2100 Bioanalyzer. cDNA libraries were sequenced in an Illumina NextSeq 6000 sequencer aiming for 240 billion 150 bp long paired-end reads.

### 2. Single-cell RNA-seq data process and analysis

Raw transcriptomic datasets of adult cochlear and vestibular HCs from scRNA-seq have been deposited to GEO (<u>GSE283534</u>). The FASTQ files were mapped to mm10 reference genome to generate the single-cell expression matrices following the CellRanger count pipeline (version 6.1.2). The Cellranger output data was then processed with the Seurat package (version 4.3.0) in R (version 4.1.3).

Genes expressed in at least ten cells were included in the analysis. Cells with numbers of expressed genes <200 or >3,000 and cells with numbers of unique molecular identifiers (UMI) > 15,000 were filtered out. Cells with >20% mitochondrial genes were also excluded from the analysis.

The gene expression data from individual samples were converted into a natural logarithm and normalized under the same condition. Data from six cochlear and four vestibular replicates were integrated separately based on the anchors identified from the top 2,000 highly variable genes of individual normalized expression matrices. The Shared Nearest Neighbor graph method can calculate the neighborhood overlap (Jaccard index) between every cell and its nearest neighbors, which was used for cluster determination at a resolution of 0.6 on PCA-reduced expression data for the top 30 principal components.

Clustering results for cochlear and vestibular datasets were visualized separately using t-distributed stochastic neighbor embedding (t-SNE). Cluster annotations were initially produced using SingleR and then corrected where appropriate based on well-known cellular markers for cochlear and vestibular cells.

Entrez Gene, HGNC, OMIM, and Ensembl database were used for verification, reference, and analyses. Online Databases of Ciliogenics (https://ciliogenics.com/), CiliaCarta (https://ngdc.cncb.ac.cn/databasecommons/database/id/6383), and Primary Cilium Proteome (https://esbl.nhlbi.nih.gov/Databases/CiliumProteome/) were also used for reference.

GO analysis in Fig. S1 was performed using ShinyGO 0.82 (Ge et al., 2020). The Venn diagrams in Fig. 5B and Fig. S3 were generated using Venny 2.1 (https://bioinfogp.cnb.csic.es/tools/venny/), except for the sperm Venn diagram in Fig. S3E, which was plotted using nVenn (https://degradome.uniovi.es/cgi-bin/nVenn/nvenn.cgi). Fig. 5C was created with BioRender.com and further modified using Adobe Photoshop.

### 3. Immunocytochemistry

Inner ears were fixed overnight with 4% paraformaldehyde at 4°C. Cochlear and vestibular sensory epithelia were dissected out. After several washes with PBS, the tissue was blocked for 1 hour with 0.25% normal goat serum in PBS containing Triton-X-100 (0.01%) and goat serum (10%). Primary antibodies against DNM1 (AB3458, Abcam), SLC7A14 (HPA045929, Sigma), TJAP1 (NBP1-80902, Novus), FOXJ1 (14-9965-82, ThermoFisher), CCDC39 (HPA035364, Sigma), CCDC40 (PA5-54653, ThermoFisher), DNAH5 (31079-1-AP, ThermoFisher), DNAH6 (HPA036391, Sigma), TEKT1 (HPA044444, Millipore Sigma), CLUAP1 (PA5-83710, Thermo Fisher), IFT172 (28441-1-AP) and acetylated tubulin (T6793, Sigma) were incubated with the tissues for 12 hours at 4°C. After washes with PBS, secondary antibody (1:500) (Alexa fluor molecular probe 488 or 555; Invitrogen) was added and incubated for 2 hours at room temperature. Alexa Fluro^TM^ 405 or 488 phalloidin (A30104 or A12379, Invitrogen) was used to label stereocilia bundles. Tissues were washed with PBS and mounted on glass microscopy slides with antifade solution (5 ml PBS, 5 ml glycerol, 0.1 g *n*-Propyl gallate). Images were captured using Nikon TI-2 Spinning Disk or Zeiss LSM 980 Inverted confocal microscope.

### 4. Single molecule fluorescence in situ hybridization

Single molecule fluorescence *in situ* hybridization was used to validate the expression of 15 genes in 10 µm-thin sections. Samples were prepared in formalin-fixed paraffin embedded tissue. Probes for 18 genes were purchased from ACD. These genes include *Adam11*, (580971), *Aqp11* (803751), *C1ql1* (465081), *Cdh23* (567261-C2), *Chrna10* (818521), *Cib2* (846681), *Cib3* (1105771), *Dnm3* (451841), *Ikzf2* (500001), *Kcnq4* (707481), *Otof* (485678), *Pcdh20* (467491), *Slc7a14* (544781), *Tmc1* (Cat No: 520911-C2) and *Tbx2* (448991-C2). Methods for the RNAscope^TM^ 2.5 HD Assay from Advanced Cell Diagnostics were followed.

### 5. Electron microscopy

The mouse inner ears were fixed with 4% paraformaldehyde and 2.5% glutaraldehyde in 0.1 M sodium cacodylate buffer (pH 7.4) with 2 mM CaCl_2_. The bullfrog crista ampullaris was fixed with 3% paraformaldehyde, 2% glutaraldehyde, 2% tannic acid and 0.5% calcium chloride in 0.1 M sodium cacodylate buffer (pH 6.8) with 0.1 mM CaCl_2_ (Kachar et al., 1990). The tissues were then post-fixed for 1 hour with 1% OsO_4_ in 0.1 M sodium cacodylate buffer and washed. For SEM, cochleae and vestibular tissues were dehydrated via an ethanol series, critical point dried from CO_2_, and sputter-coated with platinum. The morphology of the HC stereocilia bundle was examined in a FEI Quanta 200 scanning electron microscope and photographed. For TEM, the bullfrog crista ampullaris was embedded in plastic (Epon 812 Epoxy Resin) after dehydration via an ethanol series. 70-nm thin sections were cut with a diamond knife and collected on 300-mesh grids. The thin section was post-stained with 3% uranyl acetate for 15 min and 1.5% lead citrate for 3 min. The preparations were examined in an electron microscope (JEOL 100CX) and photographed.

### 6. Measurements of ciliary motion

Bullfrogs were anesthetized with 20 µg/g of 3-aminobenzoic acid ethyl ester and decapitated. The saccular macula was carefully dissected out under cooled frog’s Ringer solution. The otoliths were gently removed with forceps in fresh Ringer solution(Jaeger et al., 1994). The saccular macula was placed under an Olympus upright microscope and kinocilia motility was visualized using DIC and 60x objective and captured with a camera.

The Eustachian tube was dissected out from CBA/J mice and sectioned along its longitudinal length. The preparation was bathed in L-15 medium (Invitrogen), containing 136 mM NaCl, 5.8 mM NaH_2_PO_4_, 5.4 mM KCl, 1.4 mM CaCl_2_, 0.9 mM MgCl_2_, 0.4 mM MgSO_4_, and 10 mM HEPES-NaOH (pH 7.4, 300 mmol/l) in an experimental chamber mounted on the stage of a Leica upright microscope. Crista ampullaris was also dissected from 10-week-old CBA/J mice and bathed in L-15 medium. The tissue was attached to the bottom of the chamber by the weight of two thin platinum rods (0.5 mm in diameter). The tissue was mounted with the cilia or hair bundles facing upward toward the water-immersion objective. The cilia and hair bundles were imaged using a 63x water immersion objective (Leica) and magnified by an additional 10x relay lens. Ciliary motion was measured and calibrated by a photodiode-based measurement system mounted on the Leica upright microscope (Jia and He, 2005). The magnified image of the hair bundle was projected onto a photodiode through a rectangular slit. The image was positioned to one side of the silt with 50% of the rectangular slit being covered by the magnified image of the bundle. Cilia motion modulated the light influx to the photodiode. The photocurrent response was calibrated to displacement units by moving the slit a fixed distance (0.5 μm) with the image of the cell in front of the photodiode. After amplification, the photocurrent signal was low pass-filtered by an antialiasing filter before being digitized by a 16-bit A/D board (Digidata 1550A; Molecular Devices). The motile responses were low-pass filtered at 250 Hz and digitized at 1 kHz. Ciliary motion was acquired in a two-second window for each trial and 20 trials were captured for each cell in one recording. The power spectrum of the response was averaged and analyzed in the frequency domain using Clampfit software (version 10, Molecular Devices). The experiments were performed at room temperature (22±2°C).

### 7. Predicted model of the 96-nm modular repeat in adult vestibular kinocilia

The 96-nm repeat structures of the human respiratory axoneme (PDB: 8J07) and bovine sperm flagellum (PDB: 9FQR) were used as structural references to model the vestibular kinocilia axoneme. Candidate vestibular HC genes identified through transcriptomic profiling were annotated and mapped to their respective axonemal compartments based on known or predicted protein localizations within these reference structures. The 8J07 model lacked atomic modeling for the full-length RS3 (Zhao et al., 2025), despite its presence in the corresponding cryo-EM density map (EMD-35888), due to the uncharacterized proteome of RS3 in the human respiratory system. To address this, RS3 components from the sperm axoneme structure (PDB: 9FQR) excluding sperm-specific proteins were extracted, fitted into the EMD-35888 density using ChimeraX’s “fit-to-map” tool, and overlaid onto the 8J07 model to complete the RS3 architecture. All structural visualization and model integration were performed using UCSF ChimeraX v1.6. (Pettersen et al., 2021). In the models, DNAH5 and DNAH9 (present in our HC data) occupy the ODA region in human respiratory cilia, while DNAH8 and DNAH17 occupy the same region in sperm (but are not expressed in our dataset). Since they correspond structurally, we did not mark them as missing in the sperm-based kinocilia model.

## Acknowledgments

We acknowledge the use of the Auditory and Vestibular Technology (AVT) Core of Translational Hearing Research Center at Creighton University for high-resolution confocal imaging and library preparation (10x Genomics), and the University of Nebraska DNA Sequencing Core Facility for scRNA-seq. The AVT core at Creighton University receives partial support from NIH grant 1P20GM139762-01 from NIGMS. The University of Nebraska DNA Sequencing Core receives partial support from the NCRR (RR018788). Scanning electron microscope was acquired through a Nebraska EPSCoR MRI award (Creighton-Department of Chemistry & Biochemistry) and was wholly funded by Nebraska EPSCoR. This research was supported by the NIDCD/NIH Intramural Research Program funds Z01-DC000002 to BK and AT and utilized the computational resources of the NIH HPC Biowulf cluster (http://hpc.nih.gov).

## Funding

National Institutes of Health grant IRP funds Z01-DC000002 from NIDCD (BK and AT) National Institutes of Health grant R01 DC016807 from NIDCD (DH)

## Author contributions

D.H., B.K., and A.T. conceived the project. Z.X., H.L., S.T., Y.L., and D.H. collected the inner ear tissues and prepared the library for scRNA-seq. Z.X., A.T., S.K., C.B., J.Z., L.T., and D.H. performed the bioinformatics analyses. A.T. conducted structural modeling. A.T., H.L., S.K., D.H., and B.K. carried out morphological and physiological experiments. A.T., Z.X., S.K., C.B., B.K., and D.H. contributed to the preparation and editing of the manuscript and figures.

## Data and materials availability

Original scRNA-seq data are available (<u>GSE283534</u>). Data are visualized in the main text and the supplemental materials.

## Supplemental files include

**Fig. S1:** GO analysis of enriched genes in cellular localization and biological processes primarily related to cilia organization, assembly, maintenance, intracellular transport, and microtubule dynamics particularly associated with motile cilia

**Fig. S2:** Primary cilia-related genes in four HC types

**Fig. S3:** Comparison of HC transcriptomes with multi-tissue proteomics datasets

**Fig. S4:** ATAC-seq data indicating enhanced accessibility of motile-cilia machinery-associated gene loci in adult vestibular HCs

**Fig. S5:** Microtubule inner proteins and microtubule-associated proteins absent from mouse HC scRNA-seq data

**Table S1:** Transcriptomes of four HC types

**Table S2:** Genes/Gene products candidates associated with axonemal components in vestibular kinocilia derived from the 96-nm repeat structures of the human respiratory axoneme (PDB: 8J07). Missing genes in mouse vestibular hair cells are in red

**Videos S1 and S2**: Bullfrog kinocilia motility

**Videos S3 and S4:** Predicted structures models of molecular composition of the vestibular kinocilium based on the structural frameworks derived from human (video S3) respiratory (PDB: 8J07) (Walton et al., 2023) and bovine (video S4) sperm (PDB: 9FQR)(Leung et al., 2025) axonemes.

## Notes

https://www.ncbi.nlm.nih.gov/geo/query/acc.cgi?acc=GSE283534

### Competing Interest Statement

The authors have declared no competing interest.

### Summary of Updates

We have revised the manuscript based on comments from the reviewers. Figures 2, 5, 6, and 7 are revised. We have also added Supplemental Table 2 and two Supplemental Videos (Video 3 and Video 4). We have made changes in results and discussion.

## REFERENCE

1. Andersen JS, Vijayakumaran A, Godbehere C, Lorentzen E, Mennella V, Schou KB. 2024. Uncovering structural themes across cilia microtubule inner proteins with implications for human cilia function. Nature Communications 15:2687. DOI: 10.1038/s41467-024-46737-3

2. Anvarian Z, Mykytyn K, Mukhopadhyay S, Pedersen LB, Christensen ST. 2019. Cellular signalling by primary cilia in development, organ function and disease. Nature Reviews Nephrology 15:199–219. DOI: 10.1038/s41581-019-0116-9

3. Assad JA, Shepherd GMG, Corey DP. 1991. Tip-link integrity and mechanical transduction in vertebrate hair cells. Neuron 7:985–994. DOI: 10.1016/0896-6273(91)90343-X, PMID: 1764247

4. Baird RA. 1994. Comparative transduction mechanisms of hair cells in the bullfrog utriculus. II. Sensitivity and response dynamics to hair bundle displacement. Journal of Neurophysiology 71:685–705. DOI: 10.1152/jn.1994.71.2.685

5. Barta CL, Liu H, Chen L, Giffen KP, Li Y, Kramer KL, Beisel KW, He DZ. 2018. RNA-seq transcriptomic analysis of adult zebrafish inner ear hair cells. Scientific Data 5:180005. DOI: 10.1038/sdata.2018.5

6. Becker-Heck A, Zohn IE, Okabe N, Pollock A, Lenhart KB, Sullivan-Brown J, McSheene J, Loges NT, Olbrich H, Haeffner K, Fliegauf M, Horvath J, Reinhardt R, Nielsen KG, Marthin JK, Baktai G, Anderson KV, Geisler R, Niswander L, Omran H, Burdine RD. 2011. The coiled-coil domain containing protein CCDC40 is essential for motile cilia function and left-right axis formation. Nature Genetics 43:79–84. DOI: 10.1038/ng.727

7. Benser ME, Marquis RE, Hudspeth AJ. 1996. Rapid, Active Hair Bundle Movements in Hair Cells from the Bullfrog’s Sacculus. Journal of Neuroscience 16:5629–5643. DOI: 10.1523/JNEUROSCI.16-18-05629.1996, PMID: 8795619

8. Brody SL, Pan J, Huang T, Xu J, Xu H, Koenitizer JR, Brennan SK, Nanjundappa R, Saba TG, Rumman N, Berical A, Hawkins FJ, Wang X, Zhang R, Mahjoub MR, Horani A, Dutcher SK. 2025. Undocking of an extensive ciliary network induces proteostasis and cell fate switching resulting in severe primary ciliary dyskinesia. Science Translational Medicine 17:eadp5173. DOI: 10.1126/scitranslmed.adp5173

9. Brown A, Leung MR, Zeev-Ben-Mordehai T, Zhang R. 2025. TRiC Is a Structural Component of Mammalian Sperm Axonemes. Cytoskeleton (Hoboken, N.J.). DOI: 10.1002/cm.22005, PMID: 39927553

10. Brownell WE, Bader CR, Bertrand D, de Ribaupierre Y. 1985. Evoked mechanical responses of isolated cochlear outer hair cells. Science (New York, N.Y.) 227:194–196. DOI: 10.1126/science.3966153, PMID: 3966153

11. Burns J, Stone J. 2017. Development and regeneration of vestibular hair cells in mammals. Seminars in cell & developmental biology 65:96–105. DOI: 10.1016/j.semcdb.2016.11.001, PMID: 27864084

12. Burns JC, Kelly MC, Hoa M, Morell RJ, Kelley MW. 2015. Single-cell RNA-Seq resolves cellular complexity in sensory organs from the neonatal inner ear. Nature Communications 6:8557. DOI: 10.1038/ncomms9557

13. Chen Z, Shiozaki M, Haas KM, Skinner WM, Zhao S, Guo C, Polacco BJ, Yu Z, Krogan NJ, Lishko PV, Kaake RM, Vale RD, Agard DA. 2023. De novo protein identification in mammalian sperm using in situ cryoelectron tomography and AlphaFold2 docking. Cell 186:5041–5053.e19. DOI: 10.1016/j.cell.2023.09.017, PMID: 37865089

14. Choksi SP, Lauter G, Swoboda P, Roy S. 2014. Switching on cilia: transcriptional networks regulating ciliogenesis. Development 141:1427–1441. DOI: 10.1242/dev.074666

15. Crawford AC, Fettiplace R. 1985. The mechanical properties of ciliary bundles of turtle cochlear hair cells. The Journal of Physiology 364:359–379. DOI: 10.1113/jphysiol.1985.sp015750

16. Dallos P. 1992. The active cochlea. Journal of Neuroscience 12:4575–4585. DOI: 10.1523/JNEUROSCI.12-12-04575.1992, PMID: 1464757

17. Dallos P, Wu X, Cheatham MA, Gao J, Zheng J, Anderson CT, Jia S, Wang X, Cheng WHY, Sengupta S, He DZZ, Zuo J. 2008. Prestin-Based Outer Hair Cell Motility Is Necessary for Mammalian Cochlear Amplification. Neuron 58:333–339. DOI: 10.1016/j.neuron.2008.02.028

18. Dam TJP van, Kennedy J, Lee R van der, Vrieze E de, Wunderlich KA, Rix S, Dougherty GW, Lambacher NJ, Li C, Jensen VL, Leroux MR, Hjeij R, Horn N, Texier Y, Wissinger Y, Reeuwijk J van, Wheway G, Knapp B, Scheel JF, Franco B, Mans DA, Wijk E van, Képès F, Slaats GG, Toedt G, Kremer H, Omran H, Szymanska K, Koutroumpas K, Ueffing M, Nguyen T-MT, Letteboer SJF, Oud MM, Beersum SEC van, Schmidts M, Beales PL, Lu Q, Giles RH, Szklarczyk R, Russell RB, Gibson TJ, Johnson CA, Blacque OE, Wolfrum U, Boldt K, Roepman R, Hernandez-Hernandez V, Huynen MA. 2019. CiliaCarta: An integrated and validated compendium of ciliary genes. PLOS ONE 14:e0216705. DOI: 10.1371/journal.pone.0216705

19. Denk W, Webb WW. 1992. Forward and reverse transduction at the limit of sensitivity studied by correlating electrical and mechanical fluctuations in frog saccular hair cells. Hearing Research 60:89–102. DOI: 10.1016/0378-5955(92)90062-R

20. Eatock RA, Rüsch A, Lysakowski A, Saeki M. 1998. Hair Cells in Mammalian Utricles. Otolaryngology–Head and Neck Surgery 119:172–181. DOI: 10.1016/S0194-5998(98)70052-X

21. Eatock RA, Songer JE. 2011. Vestibular Hair Cells and Afferents: Two Channels for Head Motion Signals. Annual Review of Neuroscience 34:501–534. DOI: 10.1146/annurev-neuro-061010-113710

22. Erickson T, Biggers WP, Williams K, Butland SE, Venuto A. 2023. Regionalized Protein Localization Domains in the Zebrafish Hair Cell Kinocilium. Journal of Developmental Biology 11:28. DOI: 10.3390/jdb11020028

23. Fawcett DW, Ito S. 1965. The fine structure of bat spermatozoa. American Journal of Anatomy 116:567–609. DOI: 10.1002/aja.1001160306

24. Fettiplace R. 2017. Hair Cell Transduction, Tuning, and Synaptic Transmission in the Mammalian Cochlea. Comprehensive Physiology 7:1197–1227. DOI: 10.1002/cphy.c160049, PMID: 28915323

25. Flock Å, Flock B, Murray E. 1977. Studies on the Sensory Hairs of Receptor Cells in the Inner Ear. Acta Oto-Laryngologica 83:85–91. DOI: 10.3109/00016487709128817, PMID: 842331

26. Ge SX, Jung D, Yao R. 2020. ShinyGO: a graphical gene-set enrichment tool for animals and plants. Bioinformatics 36:2628–2629. DOI: 10.1093/bioinformatics/btz931

27. Giese APJ, Tang Y-Q, Sinha GP, Bowl MR, Goldring AC, Parker A, Freeman MJ, Brown SDM, Riazuddin S, Fettiplace R, Schafer WR, Frolenkov GI, Ahmed ZM. 2017. CIB2 interacts with TMC1 and TMC2 and is essential for mechanotransduction in auditory hair cells. Nature Communications 8:43. DOI: 10.1038/s41467-017-00061-1

28. Gillespie PG, Müller U. 2009. Mechanotransduction by Hair Cells: Models, Molecules, and Mechanisms. Cell 139:33–44. DOI: 10.1016/j.cell.2009.09.010, PMID: 19804752

29. Gui M, Farley H, Anujan P, Anderson JR, Maxwell DW, Whitchurch JB, Botsch JJ, Qiu T, Meleppattu S, Singh SK, Zhang Q, Thompson J, Lucas JS, Bingle CD, Norris DP, Roy S, Brown A. 2021. *De novo* identification of mammalian ciliary motility proteins using cryo-EM. Cell 184:5791–5806.e19. DOI: 10.1016/j.cell.2021.10.007

30. Hansen JN, Sun H, Kahnert K, Westenius E, Johannesson A, Tzavlaki K, Winsnes C, Pohjanen E, Fall J, Navarro FB, Bäckström A, Lindskog C, Johansson F, Von Feilitzen K, Vega AD, Uhlén M, Casals AM, Mahdessian D, Lindstrand A, Axelsson U, Lundberg E. 2024. Intrinsic Diversity In Primary Cilia Revealed Through Spatial Proteomics. DOI: 10.1101/2024.10.20.619273

31. Hansen JN, Sun H, Kahnert K, Westenius E, Johannesson A, Villegas C, Le T, Tzavlaki K, Winsnes C, Pohjanen E, Mäkiniemi A, Fall J, Navarro FB, Bäckström A, Lindskog C, Johansson F, Feilitzen K von, Delgado-Vega AM, Casals AM, Mahdessian D, Uhlén M, Sheu S-H, Lindstrand A, Axelsson U, Lundberg E. 2025. Intrinsic heterogeneity of primary cilia revealed through spatial proteomics. Cell 188:6804–6824.e16. DOI: 10.1016/j.cell.2025.08.039, PMID: 41005307

32. He DZZ, Dallos P. 1999. Development of Acetylcholine-Induced Responses in Neonatal Gerbil Outer Hair Cells. Journal of Neurophysiology 81:1162–1170. DOI: 10.1152/jn.1999.81.3.1162

33. Hong J, Lee C, Papoulas O, Pan J, Takagishi M, Manzi N, Dickinson DJ, Horani A, Brody SL, Marcotte E, Park TJ, Wallingford JB. 2025. Molecular organization of the distal tip of vertebrate motile cilia. DOI: 10.1101/2025.02.19.639145

34. Howard J, Hudspeth AJ. 1987. Mechanical relaxation of the hair bundle mediates adaptation in mechanoelectrical transduction by the bullfrog’s saccular hair cell. Proceedings of the National Academy of Sciences 84:3064–3068. DOI: 10.1073/pnas.84.9.3064

35. Hudspeth A. 1997. Mechanical amplification of stimuli by hair cells. Current Opinion in Neurobiology 7:480–486. DOI: 10.1016/s0959-4388(97)80026-8, PMID: 9287199

36. Jaeger RG, Fex J, Kachar B. 1994. Structural basis for mechanical transduction in the frog vestibular sensory apparatus: II. The role of microtubules in the organization of the cuticular plate. Hearing Research 77:207–215. DOI: 10.1016/0378-5955(94)90268-2, PMID: 7928733

37. Jan TA, Eltawil Y, Ling AH, Chen L, Ellwanger DC, Heller S, Cheng AG. 2021. Spatiotemporal dynamics of inner ear sensory and non-sensory cells revealed by single-cell transcriptomics. Cell Reports 36:109358. DOI: 10.1016/j.celrep.2021.109358, PMID: 34260939

38. Jen H-I, Hill MC, Tao L, Sheng K, Cao W, Zhang H, Yu HV, Llamas J, Zong C, Martin JF, Segil N, Groves AK. 2019. Transcriptomic and epigenetic regulation of hair cell regeneration in the mouse utricle and its potentiation by Atoh1. eLife 8:e44328. DOI: 10.7554/eLife.44328

39. Jia S, Dallos P, He DZZ. 2007. Mechanoelectric Transduction of Adult Inner Hair Cells. Journal of Neuroscience 27:1006–1014. DOI: 10.1523/JNEUROSCI.5452-06.2007, PMID: 17267554

40. Jia S, He DZZ. 2005. Motility-associated hair-bundle motion in mammalian outer hair cells. Nature Neuroscience 8:1028–1034. DOI: 10.1038/nn1509

41. Kachar B, Brownell WE, Altschuler R, Fex J. 1986. Electrokinetic shape changes of cochlear outer hair cells. Nature 322:365–368. DOI: 10.1038/322365a0

42. Kachar B, Parakkal M, Fex J. 1990. Structural basis for mechanical transduction in the frog vestibular sensory apparatus: I. The otolithic membrane. Hearing Research 45:179–190. DOI: 10.1016/0378-5955(90)90119-A

43. Kachar B, Parakkal M, Kurc M, Zhao Y, Gillespie PG. 2000. High-resolution structure of hair-cell tip links. Proceedings of the National Academy of Sciences 97:13336–13341. DOI: 10.1073/pnas.97.24.13336

44. Kikuchi T, Takasaka T, Tonosaki A, Watanabe H. 1989. Fine structure of guinea pig vestibular kinocilium. Acta Oto-Laryngologica 108:26–30. DOI: 10.3109/00016488909107388, PMID: 2527457

45. Kindt KS, Finch G, Nicolson T. 2012. Kinocilia Mediate Mechanosensitivity in Developing Zebrafish Hair Cells. Developmental Cell 23:329–341. DOI: 10.1016/j.devcel.2012.05.022

46. Lagziel A, Ahmed ZM, Schultz JM, Morell RJ, Belyantseva IA, Friedman TB. 2005. Spatiotemporal pattern and isoforms of cadherin 23 in wild type and *waltzer* mice during inner ear hair cell development. Developmental Biology 280:295–306. DOI: 10.1016/j.ydbio.2005.01.015

47. Lechtreck K-F, Gould TJ, Witman GB. 2013. Flagellar central pair assembly in Chlamydomonas reinhardtii. Cilia 2:15. DOI: 10.1186/2046-2530-2-15

48. Legal T, Joachimiak E, Parra M, Peng W, Tam A, Black C, Valente-Paterno M, Brouhard G, Gaertig J, Wloga D, Bui KH. 2024. Structure of the ciliary tip central pair reveals the unique role of the microtubule-seam binding protein SPEF1. DOI: 10.1101/2024.12.02.626492

49. Legal T, Parra M, Tong M, Black CS, Joachimiak E, Valente-Paterno M, Lechtreck K, Gaertig J, Bui KH. 2023. CEP104/FAP256 and associated cap complex maintain stability of the ciliary tip. Journal of Cell Biology 222:e202301129. DOI: 10.1083/jcb.202301129

50. Leibovici M, Verpy E, Goodyear RJ, Zwaenepoel I, Blanchard S, Lainé S, Richardson GP, Petit C. 2005. Initial characterization of kinocilin, a protein of the hair cell kinocilium. Hearing Research 203:144–153. DOI: 10.1016/j.heares.2004.12.002

51. Leung MR, Sun C, Zeng J, Anderson JR, Niu Q, Huang W, Noteborn WEM, Brown A, Zeev-Ben-Mordehai T, Zhang R. 2025. Structural diversity of axonemes across mammalian motile cilia. Nature 1–8. DOI: 10.1038/s41586-024-08337-5

52. Li A, Xue J, Peterson EH. 2008. Architecture of the Mouse Utricle: Macular Organization and Hair Bundle Heights. Journal of Neurophysiology 99:718–733. DOI: 10.1152/jn.00831.2007

53. Li Y, Liu H, Giffen KP, Chen L, Beisel KW, He DZZ. 2018. Transcriptomes of cochlear inner and outer hair cells from adult mice. Scientific Data 5:180199. DOI: 10.1038/sdata.2018.199

54. Liberman MC, Gao J, He DZZ, Wu X, Jia S, Zuo J. 2002. Prestin is required for electromotility of the outer hair cell and for the cochlear amplifier. 419:5.

55. Liu H, Pecka JL, Zhang Q, Soukup GA, Beisel KW, He DZZ. 2014. Characterization of Transcriptomes of Cochlear Inner and Outer Hair Cells. Journal of Neuroscience 34:11085–11095. DOI: 10.1523/JNEUROSCI.1690-14.2014, PMID: 25122905

56. Lysakowski A, Goldberg JM. 1997. A regional ultrastructural analysis of the cellular and synaptic architecture in the chinchilla cristae ampullares. Journal of Comparative Neurology 389:419–443. DOI: 10.1002/(SICI)1096-9861(19971222)389:3%253C419::AID-CNE5%253E3.0.CO;2-3

57. Ma Y, He J, Li S, Yao D, Huang C, Wu J, Lei M. 2023. Structural insight into the intraflagellar transport complex IFT-A and its assembly in the anterograde IFT train. Nature Communications 14:1506. DOI: 10.1038/s41467-023-37208-2

58. Martin P, Bozovic D, Choe Y, Hudspeth AJ. 2003. Spontaneous Oscillation by Hair Bundles of the Bullfrog’s Sacculus. Journal of Neuroscience 23:4533–4548. DOI: 10.1523/JNEUROSCI.23-11-04533.2003, PMID: 12805294

59. McInturff S, Burns JC, Kelley MW. 2018. Characterization of spatial and temporal development of Type I and Type II hair cells in the mouse utricle using new cell-type-specific markers. Biology Open 7:bio038083. DOI: 10.1242/bio.038083

60. Meng G-Q, Wang Y, Luo C, Tan Y-M, Li Y, Tan C, Tu C, Zhang Q-J, Hu L, Zhang H, Meng L-L, Liu C-Y, Deng L, Lu G-X, Lin G, Du J, Tan Y-Q, Sha Y, Wang L, He W-B. 2024. Bi-allelic variants in DNAH3 cause male infertility with asthenoteratozoospermia in humans and mice. Human Reproduction Open 2024:hoae003. DOI: 10.1093/hropen/hoae003, PMID: 38312775

61. Meng X, Li L, Lin C, Zhu Y, Feng Y, Zhou X, Tong Y, Wang S, Yin G, Liu R, Sun F, Yan X, Zhu X, Cong Y. 2024. Chaperonin TRiC bridges radial spokes for folding proteins locally translated in mammalian sperm flagella. DOI: 10.1101/2024.11.19.624414

62. Merveille A-C, Davis EE, Becker-Heck A, Legendre M, Amirav I, Bataille G, Belmont J, Beydon N, Billen F, Clément A, Clercx C, Coste A, Crosbie R, de Blic J, Deleuze S, Duquesnoy P, Escalier D, Escudier E, Fliegauf M, Horvath J, Hill K, Jorissen M, Just J, Kispert A, Lathrop M, Loges NT, Marthin JK, Momozawa Y, Montantin G, Nielsen KG, Olbrich H, Papon J-F, Rayet I, Roger G, Schmidts M, Tenreiro H, Towbin JA, Zelenika D, Zentgraf H, Georges M, Lequarré A-S, Katsanis N, Omran H, Amselem S. 2011. CCDC39 is required for assembly of inner dynein arms and the dynein regulatory complex and for normal ciliary motility in humans and dogs. Nature Genetics 43:72–78. DOI: 10.1038/ng.726

63. Moon K-H, Ma J-H, Min H, Koo H, Kim H, Ko HW, Bok J. 2020. Dysregulation of sonic hedgehog signaling causes hearing loss in ciliopathy mouse models. eLife 9:e56551. DOI: 10.7554/eLife.56551

64. Nagel ZD, Margulies CM, Chaim IA, McRee SK, Mazzucato P, Ahmad A, Abo RP, Butty VL, Forget AL, Samson LD. 2014. Multiplexed DNA repair assays for multiple lesions and multiple doses via transcription inhibition and transcriptional mutagenesis. Proceedings of the National Academy of Sciences 111:E1823–E1832. DOI: 10.1073/pnas.1401182111

65. Oda T, Yanagisawa H, Kamiya R, Kikkawa M. 2014. A molecular ruler determines the repeat length in eukaryotic cilia and flagella. Science 346:857–860. DOI: 10.1126/science.1260214

66. O’Donnell J, Zheng J. 2022. Vestibular Hair Cells Require CAMSAP3, a Microtubule Minus-End Regulator, for Formation of Normal Kinocilia. Frontiers in Cellular Neuroscience 16:876805. DOI: 10.3389/fncel.2022.876805, PMID: 35783105

67. Pettersen EF, Goddard TD, Huang CC, Meng EC, Couch GS, Croll TI, Morris JH, Ferrin TE. 2021. UCSF ChimeraX: Structure visualization for researchers, educators, and developers. Protein Science 30:70–82. DOI: 10.1002/pro.3943

68. Pir MS, Begar E, Yenisert F, Demirci HC, Korkmaz ME, Karaman A, Tsiropoulou S, Firat-Karalar EN, Blacque OE, Oner SS, Doluca O, Cevik S, Kaplan OI. 2024. CilioGenics: an integrated method and database for predicting novel ciliary genes. Nucleic Acids Research 52:8127–8145. DOI: 10.1093/nar/gkae554

69. Polino AJ, Sviben S, Melena I, Piston DW, Hughes JW. 2023. Scanning electron microscopy of human islet cilia. Proceedings of the National Academy of Sciences 120:e2302624120. DOI: 10.1073/pnas.2302624120

70. Pujol R, Pickett SB, Nguyen TB, Stone JS. 2014. Large basolateral processes on type II hair cells comprise a novel processing unit in mammalian vestibular organs. The Journal of comparative neurology 522:3141–3159. DOI: 10.1002/cne.23625, PMID: 24825750

71. Ranum PT, Goodwin AT, Yoshimura H, Kolbe DL, Walls WD, Koh J-Y, He DZZ, Smith RJH. 2019. Insights into the Biology of Hearing and Deafness Revealed by Single-Cell RNA Sequencing. Cell Reports 26:3160–3171.e3. DOI: 10.1016/j.celrep.2019.02.053

72. Riazuddin Saima, Belyantseva IA, Giese APJ, Lee K, Indzhykulian AA, Nandamuri SP, Yousaf R, Sinha GP, Lee S, Terrell D, Hegde RS, Ali RA, Anwar S, Andrade-Elizondo PB, Sirmaci A, Parise LV, Basit S, Wali A, Ayub M, Ansar M, Ahmad W, Khan SN, Akram J, Tekin M, Riazuddin Sheikh, Cook T, Buschbeck EK, Frolenkov GI, Leal SM, Friedman TB, Ahmed ZM. 2012. Alterations of the CIB2 calcium- and integrin-binding protein cause Usher syndrome type 1J and nonsyndromic deafness DFNB48. Nature Genetics 44:1265–1271. DOI: 10.1038/ng.2426

73. Ricci AJ, Crawford AC, Fettiplace R. 2003. Tonotopic Variation in the Conductance of the Hair Cell Mechanotransducer Channel. Neuron 40:983–990. DOI: 10.1016/S0896-6273(03)00721-9

74. Ricci AJ, Rennie KJ, Cochran SL, Kevetter GA, Correia MJ. 1997. Vestibular type I and type II hair cells. 1: Morphometric identification in the pigeon and gerbil. Journal of Vestibular Research: Equilibrium & Orientation 7:393–406. PMID: 9376913

75. Rüsch A, Thurm U. 1990. Spontaneous and electrically induced movements of ampullary kinocilia and stereovilli. Hearing Research 48:247–263. DOI: 10.1016/0378-5955(90)90065-W

76. Schwander M, Kachar B, Müller U. 2010. The cell biology of hearing. Journal of Cell Biology 190:9–20. DOI: 10.1083/jcb.201001138

77. Shi H, Wang H, Zhang C, Lu Y, Yao J, Chen Z, Xing G, Wei Q, Cao X. 2022. Mutations in OSBPL2 cause hearing loss associated with primary cilia defects via sonic hedgehog signaling. DOI: 10.1172/jci.insight.149626

78. Shin J-B, Krey JF, Hassan A, Metlagel Z, Tauscher AN, Pagana JM, Sherman NE, Jeffery ED, Spinelli KJ, Zhao H, Wilmarth PA, Choi D, David LL, Auer M, Barr-Gillespie PG. 2013. Molecular architecture of the chick vestibular hair bundle. Nature Neuroscience 16:365–374. DOI: 10.1038/nn.3312

79. Silver IA, Erecińska M. 1997. Energetic demands of the Na+/K+ ATPase in mammalian astrocytes. Glia 21:35–45. DOI: 10.1002/(SICI)1098-1136(199709)21:1%253C35::AID-GLIA4%253E3.0.CO;2-0

80. Spoon C, Grant W. 2011. Biomechanics of hair cell kinocilia: experimental measurement of kinocilium shaft stiffness and base rotational stiffness with Euler–Bernoulli and Timoshenko beam analysis. Journal of Experimental Biology 214:862–870. DOI: 10.1242/jeb.051151

81. Stooke-Vaughan GA, Huang P, Hammond KL, Schier AF, Whitfield TT. 2012. The role of hair cells, cilia and ciliary motility in otolith formation in the zebrafish otic vesicle. Development 139:1777–1787. DOI: 10.1242/dev.079947

82. Sung YH, Baek I-J, Kim YH, Gho YS, Oh SP, Lee YJ, Lee H-W. 2016. PIERCE1 is critical for specification of left-right asymmetry in mice. Scientific Reports 6:27932. DOI: 10.1038/srep27932

83. Tai L, Yin G, Huang X, Sun F, Zhu Y. 2023. In-cell structural insight into the stability of sperm microtubule doublet. Cell Discovery 9:1–19. DOI: 10.1038/s41421-023-00606-3

84. Tian X, Zhao H, Zhou J. 2023. Organization, functions, and mechanisms of the BBSome in development, ciliopathies, and beyond. eLife 12:e87623. DOI: 10.7554/eLife.87623

85. Vasquez SSV, van Dam J, Wheway G. 2021. An updated SYSCILIA gold standard (SCGSv2) of known ciliary genes, revealing the vast progress that has been made in the cilia research field. Molecular Biology of the Cell 32:br13. DOI: 10.1091/mbc.E21-05-0226

86. Walton T, Gui M, Velkova S, Fassad MR, Hirst RA, Haarman E, O’Callaghan C, Bottier M, Burgoyne T, Mitchison HM, Brown A. 2023. Axonemal structures reveal mechanoregulatory and disease mechanisms. Nature 618:625–633. DOI: 10.1038/s41586-023-06140-2

87. Wang D, Zhou J. 2021. The Kinocilia of Cochlear Hair Cells: Structures, Functions, and Diseases. Frontiers in Cell and Developmental Biology 9. DOI: 10.3389/fcell.2021.715037

88. Wang T, Ling AH, Billings SE, Hosseini DK, Vaisbuch Y, Kim GS, Atkinson PJ, Sayyid ZN, Aaron KA, Wagh D, Pham N, Scheibinger M, Zhou R, Ishiyama A, Moore LS, Maria PS, Blevins NH, Jackler RK, Alyono JC, Kveton J, Navaratnam D, Heller S, Lopez IA, Grillet N, Jan TA, Cheng AG. 2024. Single-cell transcriptomic atlas reveals increased regeneration in diseased human inner ear balance organs. Nature Communications 15:4833. DOI: 10.1038/s41467-024-48491-y

89. Wang X, Liu S, Cheng Q, Qu C, Ren R, Du H, Li N, Yan K, Wang Y, Xiong W, Xu Z. 2023. CIB2 and CIB3 Regulate Stereocilia Maintenance and Mechanoelectrical Transduction in Mouse Vestibular Hair Cells. Journal of Neuroscience 43:3219–3231. DOI: 10.1523/JNEUROSCI.1807-22.2023, PMID: 37001993

90. Wilkerson BA, Zebroski HL, Finkbeiner CR, Chitsazan AD, Beach KE, Sen N, Zhang RC, Bermingham-McDonogh O. 2021. Novel cell types and developmental lineages revealed by single-cell RNA-seq analysis of the mouse crista ampullaris. eLife 10:e60108. DOI: 10.7554/eLife.60108

91. Wu D, Freund JB, Fraser SE, Vermot J. 2011. Mechanistic Basis of Otolith Formation during Teleost Inner Ear Development. Developmental Cell 20:271–278. DOI: 10.1016/j.devcel.2010.12.006, PMID: 21316594

92. Xia X, Shimogawa MM, Wang H, Liu S, Wijono A, Langousis G, Kassem AM, Wohlschlegel JA, Hill KL, Zhou ZH. 2025. Trypanosome doublet microtubule structures reveal flagellum assembly and motility mechanisms. Science 387:eadr3314. DOI: 10.1126/science.adr3314

93. Yamamoto R, Song K, Yanagisawa H, Fox L, Yagi T, Wirschell M, Hirono M, Kamiya R, Nicastro D, Sale WS. 2013. The MIA complex is a conserved and novel dynein regulator essential for normal ciliary motility. Journal of Cell Biology 201:263–278. DOI: 10.1083/jcb.201211048

94. Yang M, Hussain HMJ, Khan M, Muhammad Z, Zhou J, Ma A, Huang X, Ye J, Chen M, Zhi A, Liu T, Khan R, Ali A, Shah W, Zeb A, Ahmad N, Zhang H, Xu B, Ma H, Shi Q, Shi B. 2024. Deficiency in a special dynein DNAH12 causes male infertility by impairing DNAH1 and DNALI1 recruitment in humans and mice. eLife 13. DOI: 10.7554/eLife.100350.1

95. Zhao Y, Song K, Tavakoli A, Gui L, Fernandez-Gonzalez A, Zhang S, Dzeja PP, Mitsialis SA, Zhang X, Nicastro D. 2025. Mouse radial spoke 3 is a metabolic and regulatory hub in cilia. Nature Structural & Molecular Biology 32:1542–1554. DOI: 10.1038/s41594-025-01594-6

96. Zheng J, Shen W, He DZZ, Long KB, Madison LD, Dallos P. 2000. Prestin is the motor protein of cochlear outer hair cells. Nature 405:149–155. DOI: 10.1038/35012009

