## Supplemental Table 2 for "The Dual Molecular Identity of Vestibular Kinocilia: Bridging Structural and Functional Traits of Primary and Motile Cilia"

**Table S2.** **Genes/ Gene products candidates associated with axonemal components in vestibular kinocilia derived from the 96-nm repeat structures of the human respiratory axoneme (PDB: 8J07). Missing genes in mouse vestibular hair cells are in red**

| Axonemal complex | Axonemal protein | Present in this study | Gene/Transcript in this study |
|---|---|---|---|
| **ODA** | DNAH5 | Yes | |
| | DNAH9 | Yes | |
| | DNAI1/Dnaic1 | Yes | Dnaic1 |
| | DNAI2/Dnaic2 | Yes | Dnaic2 |
| | DNAL1 | Yes | |
| | DYNLRB1 | Yes | |
| | DYNLRB2 | Yes | |
| | DYNLL1 (and/or DYNLL2) | Yes | |
| | DYNLT2B (and/or DYNLT2, DYNLT4) | Yes | DYT2B |
| | DYNLT1 (and/or DYNLT3) (TCTEX1) | Yes | DYLT1 |
| | NME9 | Yes | |
| **ODA-DC** | ODAD1 (CCDC114) | Yes | CCDC114 |
| | ODAD2 (ARMC4) | Yes | ARMC4 |
| | ODAD3 (CCDC151) | Yes | CCDC151 |
| | ODAD4 (TTC25) | Yes | TTC25 |
| | ODAD5 (clxn, EFCAB1) | Yes | EFCAB1 |
| **IDAf** | DNAH2 | Yes | |
| | DNAH10 | Yes | |
| | DNAI3/(WDR63) | Yes | WDR63 |
| | DNAI4/(WDR78) | Yes | WDR78 |
| | DNAI7/LAS1/CFAP94/CASC1 | Yes | CASC1 |
| | DYNLRB1 | Yes | |
| | DYNLRB2 | Yes | |
| | DYNLT2B (and/or DYNLT2, DYNLT4) | Yes | DYT2B/ Tctex1d2 |
| | DYNLT1 (and/or DYNLT3) (TCTEX1) | Yes | Dynlt3 |
| | CFAP57 | Yes | |
| | LRRC51/DFNB63 | Yes | |
| **MIA** | CFAP73 | Yes | |
| | CFAP100/CCDC37 | No | |
| **T/TH** | CFAP43 | Yes | |
| | CFAP44 | Yes | |
| **Single head-IDAs** | DNAH1(IDAd) | Yes | |
| | DNAH3(IDAc) | No | |
| | DNAH6(IDAg) | Yes | |
| | DNAH7(IDAb/e) | Yes | |
| | DNAH12 (IDAa) | No | |
| | DNALI1(IDAa) | Yes | |
| | CEN2 | Yes | CETN2 |
| | ACTA2 | No | |
| | TTC29 | Yes | |
| | ZMYND12 | Yes | |
| **N-DRC** | DRC1/CCDC164 | Yes | DRC1 |
| | DRC2/CCDC65 | Yes | CCDC65 |
| | DRC3/LRRC48 | Yes | DRC3 |
| | DRC4/GAS8 | Yes | GAS8 |
| | DRC5/TCTE1 | Yes | TCTE1 |
| | DRC7 | Yes | DRC7 |
| | DRC8/CCDC135 | Yes | EFCAB2 |
| | DRC9/IQCG | Yes | IQCG |
| | DRC10/IQCD | Yes | IQCD |
| | DRC11/IQCA | Yes | IQCA |
| | ANKEF1 | Yes | |
| **RS** | CALM1 | Yes | |
| | CYB5D1 | Yes | |
| | DNAJB13 | Yes | |
| | DYDC2 | Yes | |

| | | | |
|---|---|---|---|
| | DYNLL1 | Yes | |
| | IQUB | Yes | |
| | LRRC34 | Yes | |
| | MORN3 | No | |
| | NME5 | Yes | |
| | PPIL6 | Yes | |
| | ROPN1L | Yes | |
| | RSPH1 | Yes | |
| | RSPH3 | Yes | |
| | RSPH4A | Yes | |
| | RSPH9 | Yes | |
| | RSPH14 | Yes | |
| | SPA17 | Yes | |
| | AK7 | Yes | |
| | AK9 | Yes | |
| | AKAP14 | Yes | |
| | CATIP | Yes | |
| | LRRC23 | Yes | |
| | MDH1A | Yes | |
| | MDH1B | Yes | |
| | MORN5 | Yes | |
| | PRKACA | Yes | |
| | PRKAR1A/1B | Yes | |
| | STYXL1 | Yes | |
| | AK8 | Yes | |
| | CFAP206 | Yes | |
| | EFCAB10 | Yes | |
| | LRP2BP | Yes | |
| | RGS22 | Yes | |
| | LRRC23 | Yes | |
| | CFAP91/MAATS1 | Yes | MAATS1 |
| | CFAP69 | Yes | |
| | FAP251/WDR66 | Yes | WDR66 |

| | | | |
|---|---|---|---|
| **MIPs** | CFAP20 | Yes | |
| | CFAP45 | Yes | |
| | CFAP52 | Yes | |
| | CFAP53 | Yes | |
| | CFAP68 | No | |
| | CFAP77 | Yes | |
| | CFAP90 | No | |
| | CFAP95 | No | |
| | CFAP107 | No | |
| | CFAP126 | Yes | |
| | CFAP141 | No | |
| | CFAP161 | Yes | |
| | CFAP210/CCDC173 | Yes | CCDC173 |
| | CFAP276 | No | |
| | EFHC1 | Yes | |
| | EFHC2 | Yes | |
| | ENKUR | Yes | |
| | FAM166B//CIMIP2B | No | |
| | FAM183A/CFAP144 | No | |
| | FAM166C | No | |
| | MNS1 | Yes | |
| | NME7 | Yes | |
| | PACRG | Yes | |
| | Pierce 1 | No | |
| | Pierce 2 | No | |
| | RIBC1 | Yes | |
| | RIBC2 | Yes | |
| | SPACA9 | Yes | |
| | SPAG8 | Yes | |
| | Tektin 1/Tekt1 | Yes | Tekt1 |
| | Tektin 2/Tekt2 | Yes | Tekt2 |

| | | |
|---|---|---|
| **External coiled coils** | CCDC39 | Yes |
| | CCDC40 | Yes |
| | CCDC96 | Yes |
| | CCDC113 | Yes |
| | CFAP299 | Yes |
| | CFAP58 | Yes |
| | CCDC146 | Yes |
| | ARMH1 | Yes |
