## Supplemental Figure 1 to Figure 5 for "The Dual Molecular Identity of Vestibular Kinocilia: Bridging Structural and Functional Traits of Primary and Motile Cilia"

**Supplemental Figures for Manuscript by Xu et al:**

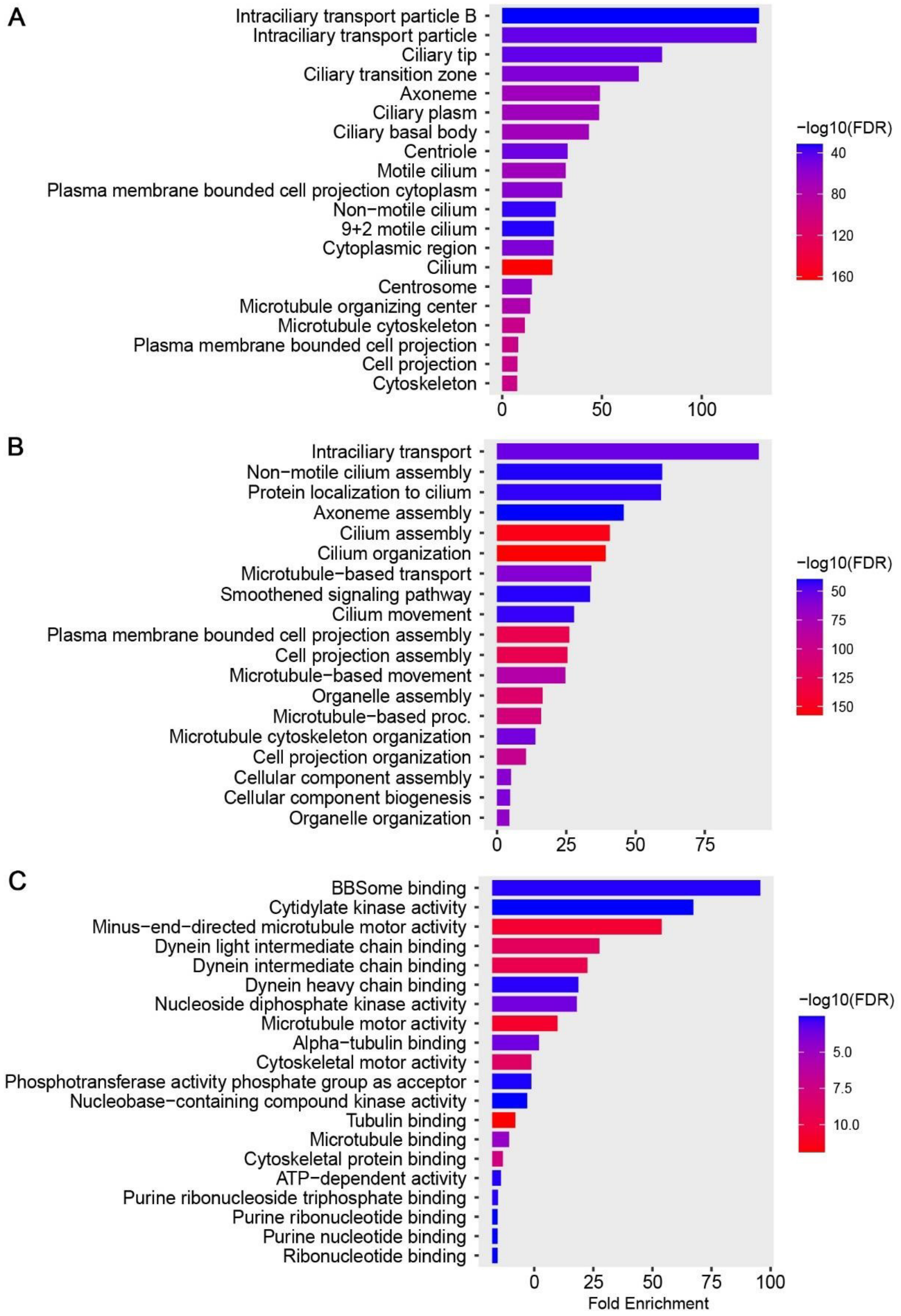


**Fig. S1:** Gene Ontology (GO) analysis of shared genes in panel 5B across multiple ciliary databases and the current study. (A) Cellular localization, (B) Molecular function, and (C) Biological processes. The analysis reveals enrichment in terms related to cilia organization, assembly, maintenance, intracellular transport, and microtubule dynamics, particularly those associated with motile cilia.

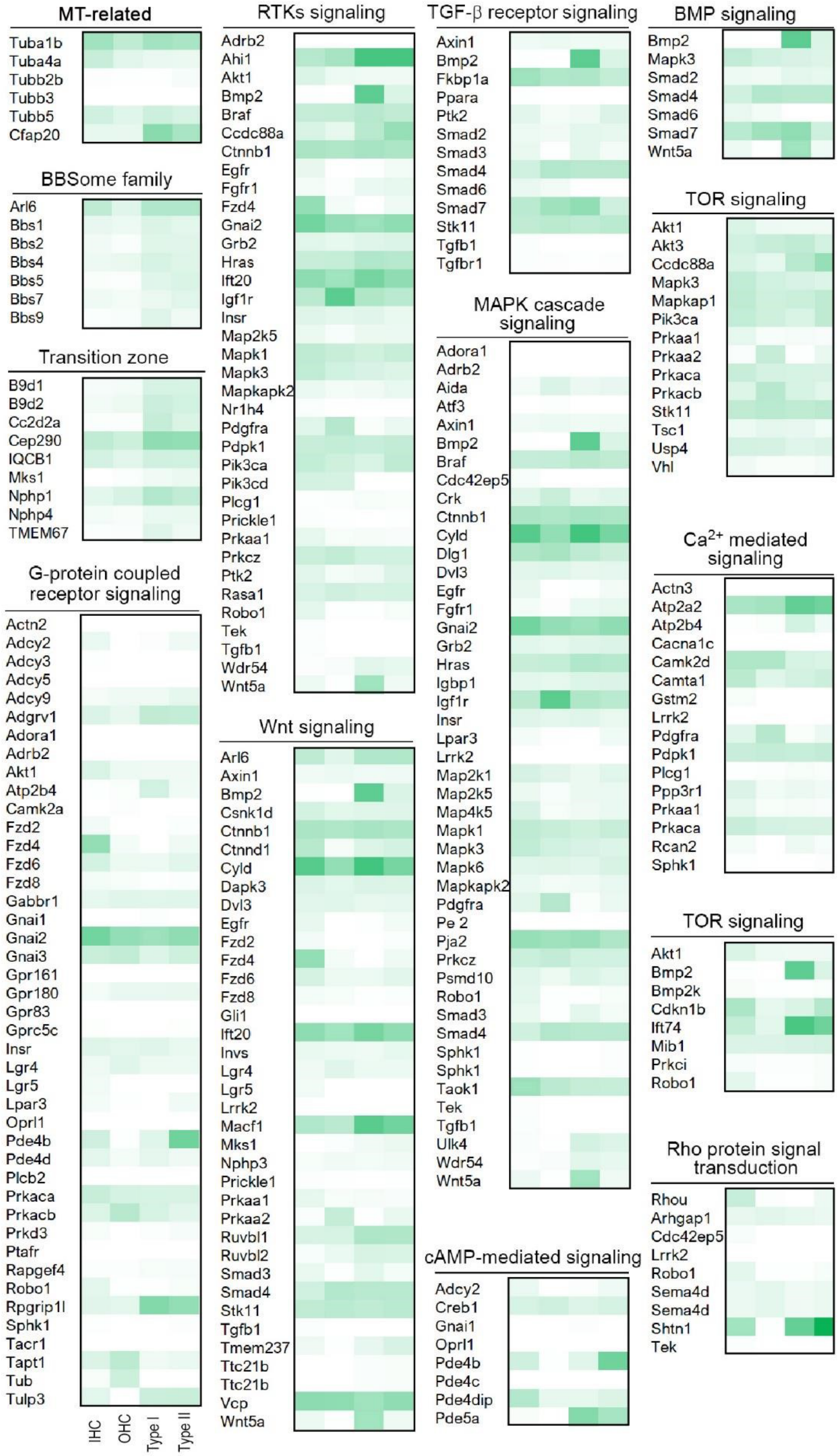


**Fig. S2**: Primary cilia-related genes in four HC subtypes. Overlap of shared genes involved in cilia maintenance, including microtubule-associated proteins, BBSome family members, and components of the transition zone and primary cilia signaling pathways associated with primary cilia and their basal body.

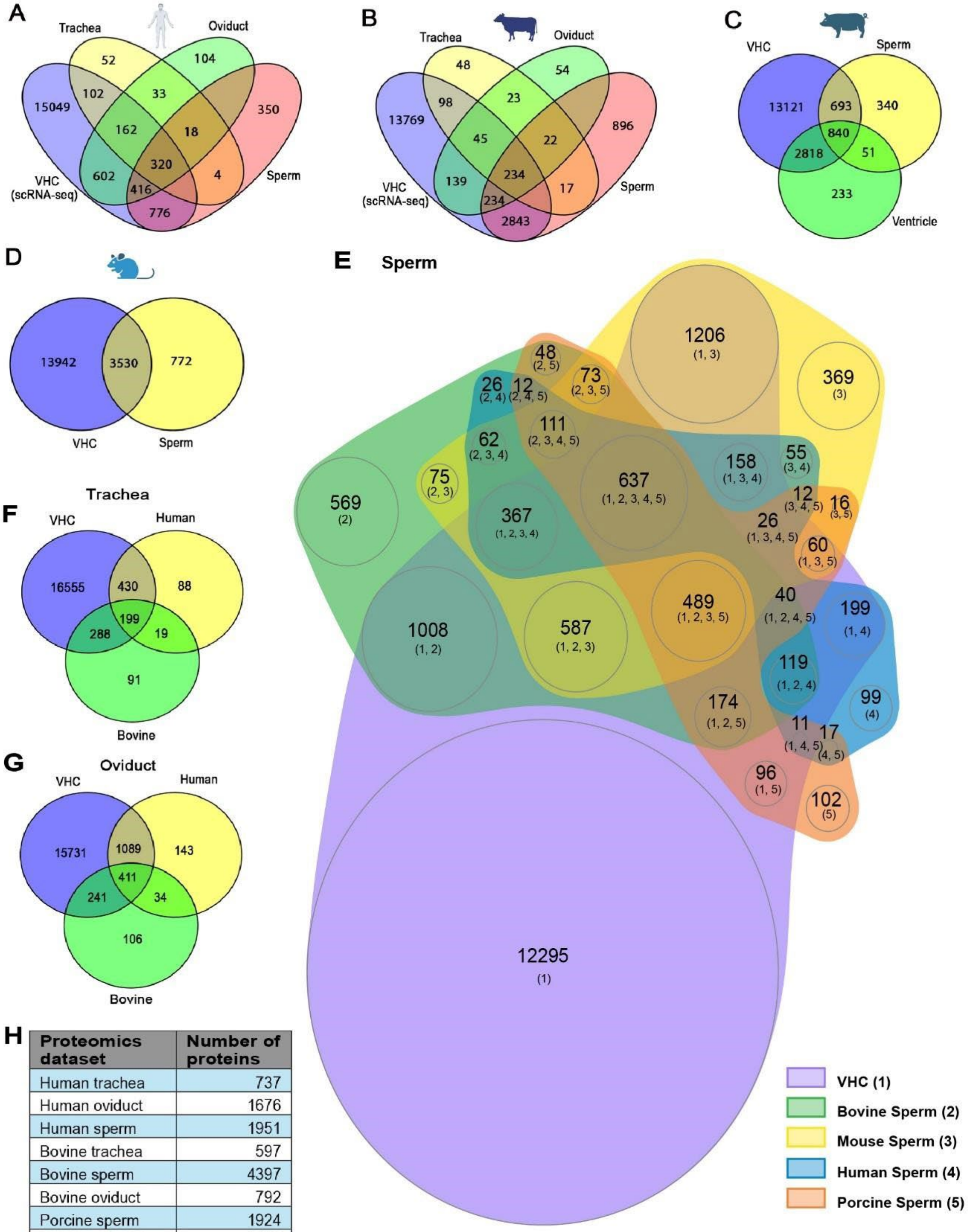


| Proteomics dataset | Number of proteins |
|---|---|
| Human trachea | 737 |
| Human oviduct | 1676 |
| Human sperm | 1951 |
| Bovine trachea | 597 |
| Bovine sperm | 4397 |
| Bovine oviduct | 792 |
| Porcine sperm | 1924 |
| Porcine ventricle | 3492 |
| Mouse sperm | 4302 |

**Fig. S3: Comparison of vestibular HC scRNA-seq data with proteomics datasets from multiple tissues and species containing motile cilia.** (A–D) Cross-species comparisons of motile cilia proteomes from different tissues, each compared with VHC scRNA-seq data: (A) Human trachea, oviduct, and sperm. (B) Bovine trachea, oviduct, and sperm. (C) Porcine sperm and ventricle. (D) Mouse sperm. (E–G) Tissue-specific comparisons across species, each compared with VHC scRNA-seq data: (E) Sperm proteomes from bovine, mouse, human, and porcine. (F) Tracheal proteomes from human and bovine. (G) Oviduct proteomes from human and bovine. (H) Number of proteins identified in each dataset included in the comparisons.

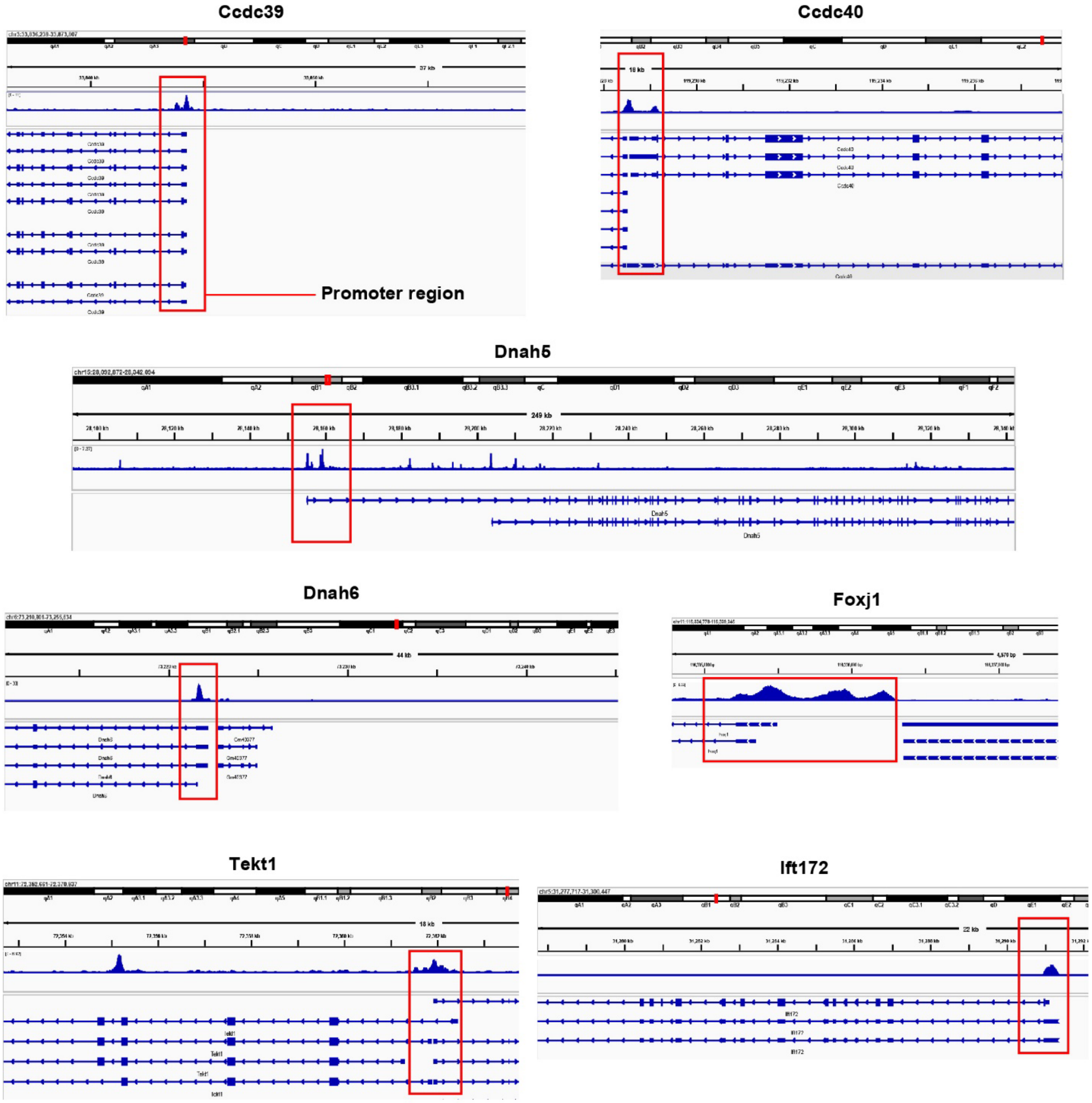


**Fig. S4:** Enhanced accessibility of motile cilia-associated gene loci in adult vestibular HCs based on published ATAC-seq data

| Axonemal complex | Missing genes | Corresponding PDB model | Effect on motility/Cilia-related disease |
|---|---|---|---|
| MIPs and MAPs | CFAP68 | 8J07/9FQR | unknown |
| | CFAP90 | 8J07/9FQR | unknown |
| | CFAP95 | 8J07/9FQR | unknown |
| | CFAP107 | 8J07/9FQR | unknown |
| | CFAP141 | 8J07/9FQR | unknown |
| | CFAP276 | 8J07/9FQR | unknown |
| | FAM166B | 8J07/9FQR | unknown |
| | FAM183A | 8J07/9FQR | unknown |
| | FAM166C | 8J07/9FQR | unknown |
| | Pierce1 | 8J07/9FQR | LR asymmetry<br>Defect on ciliary motility and laterality |
| | Pierce2 | 8J07/9FQR | LR asymmetry<br>Defect on ciliary motility and laterality |
| | SPMIP1 | 9FQR | unknown |
| | CIMIP1 | 9FQR | unknown |
| | SPMAP1 | 9FQR | unknown |
| | CFAP 144 | 9FQR | unknown |
| | CFAP 263 | 9FQR | unknown |
| | CIMAP1B | 9FQR | unknown |
| | CIMAP1C | 9FQR | unknown |
| | CIMIP3 | 9FQR | unknown |
| | CCDC105 | 9FQR | unknown |
| | DUSP21 | 9FQR | unknown |
| | EFCAB3 | 9FQR | unknown |
| | ODF3 | 9FQR | unknown |
| | SIMIP4 | 9FQR | unknown |
| | SPMIP7 | 9FQR | unknown |
| | SPMIP5 | 9FQR | unknown |
| | SPMIP6 | 9FQR | unknown |
| | SAXO3 | 9FQR | unknown |
| | TEKTIP1 | 9FQR | unknown |
| | Tektin-5 | 9FQR | unknown |
| | SPMIP8 | 9FQR | unknown |
| | TEX33 | 9FQR | unknown |
| | TEX49 | 9FQR | unknown |
| | TSSk6 | 9FQR | unknown |
| | TEX37 | 9FQR | unknown |
| | THEGL | 9FQR | unknown |
| | CFAP 96 | 9FQR | unknown |
| | CFAP 299 | 9FQR | unknown |

**Fig. S5: Microtubule inner proteins (MIPs) and microtubule-associated proteins (MAPs) absent from mouse HC scRNA-seq data.** These genes structurally resolved in the 96-nm axonemal repeats of human respiratory and/or bovine sperm flagella (PDB: 8J07, 9FQR). While most of these proteins are not known to directly regulate motility, a few such as Pierce1 and Pierce2 are linked to defects in ciliary motility and left–right (LR) asymmetry.
